# Systematic discovery of receptor-ligand biology by engineered cell entry and single-cell genomics

**DOI:** 10.1101/2021.12.13.472464

**Authors:** Bingfei Yu, Quanming Shi, Julia A. Belk, Kathryn E. Yost, Kevin R. Parker, Huang Huang, Daniel Lingwood, Mark M. Davis, Ansuman T. Satpathy, Howard Y. Chang

**Author notes:** Co-first authors (Bingfei Yu, Quanming Shi).

## Abstract

Cells communicate with each other via receptor-ligand interactions on the cell surface. Here we describe a technology for l**<u>e</u>**ntiviral-mediated cell e**<u>nt</u>**ry by **<u>e</u>**ngineered **<u>r</u>**eceptor-ligand interaction (ENTER) to decode receptor specificity. Engineered lentiviral particles displaying specific ligands deliver fluorescent proteins into target cells upon cognate receptor-ligand interaction, without genome integration or transgene transcription. We optimize ENTER to decode interactions between T cell receptor (TCR)-MHC peptides, antibody-antigen, and other receptor-ligand pairs. We develop an effective presentation strategy to capture interactions between B cell receptor (BCR) and intracellular antigen epitopes. Single-cell readout of ENTER by RNA sequencing (ENTER-seq) enables multiplexed enumeration of TCR-antigen specificities, clonality, cell type, and cell states of individual T cells. ENTER-seq of patient blood samples after CMV infection reveals the viral epitopes that drive human effector memory T cell differentiation and inter-clonal phenotypic diversity that targets the same epitope. ENTER enables systematic discovery of receptor specificity, linkage to cell fates, and cell-specific delivery of gene or protein payloads.

**HIGHLIGHTS:**

- ENTER displays ligands, deliver cargos, and records receptor specificity.
- ENTER deorphanizes antigen recognition of TCR and BCR.
- ENTER-seq maps TCR specificity, clonality and cell state in single cells.
- ENTER-seq of patient sample decodes antiviral T cell memory.

## INTRODUCTION

Advances in single-cell genomics field have provided unprecedented insights in deciphering molecular, cellular, and phenotypic heterogeneity of biological systems. High-throughput profiling at a single-cell resolution uncovers molecular information including gene expression, chromatin accessibility, histone modifications, surface protein markers, and immune receptor repertoires (Buenrostro et al., 2015; Kaya-Okur et al., 2019; Stoeckius et al., 2017; Yost et al., 2019). Integration of multiple layers of molecular information has been achieved by single-cell multiomic tools such as ECCITE-seq (transcriptome & surface protein & CRISPR guide RNA & T cell receptor (TCR)), Perturb-ATAC (CRISPR guide RNA & chromatin accessibility), and TCR-ATAC (TCR sequence & chromatin accessibility) (Mimitou et al., 2019; Rubin et al., 2019; Satpathy et al., 2018). Current single-cell tools have mainly centered on deciphering the central dogma of gene expression in a cell-intrinsic manner. However, cell-extrinsic information such as intercellular ligand-receptor interaction is poorly explored at the single-cell resolution due to the lack of experimental tools.

Cells communicate with each other through ligand-receptor interactions. Rich intercellular communications shape the molecular programs of mammalian cells to instruct specific behaviors and cell fate decisions (Armingol et al., 2021). For example, TCR on the surface of T cells can recognize and interact with the major histocompatibility complex (MHC)–antigen complexes (pMHC) on the surface of antigen-presenting cells (APCs) (Davis and Bjorkman, 1988). TCR and antibody genes undergo somatic recombination to reach a large and diverse repertoire (∼10^19 TCR alpha and beta combinations in humans), which are clonally inherited by daughter cells (Robins et al., 2009). T and B cell receptor interactions are highly specific and drive antigen-specific T and B cell expansion and differentiation. Resolving TCR-antigen interactions, especially linking antigen specificity to TCR sequences and T cell states are essential to understand how antigen recognition drives T cell fate decisions (Pai and Satpathy, 2021). Diverse approaches have been developed to decipher the antigen specificity of TCRs including: (1) Cell reporter assay to screen T cell-specific pMHCs using artificial APCs such as T-scan, SABR, T cell trogocytosis, and cytokine capturing assay (Joglekar et al., 2019; Kula et al., 2019; Lee and Meyerson, 2021; Li et al., 2019); (2) yeast display platform to screen pMHCs for recombinant TCRs (Birnbaum et al., 2012); (3) T cell based assay such as cytokine production (ELISPOT) upon antigen peptide stimulation (McCutcheon et al., 1997); (4) DNA barcoded pMHC tetramer to capture antigen specificity and TCR sequence by single cell sequencing (TetTCR-seq) (Zhang et al., 2018). Despite unique advantages for each technique, it is still challenging to rapidly screen immunogenic pMHCs for primary T cells and simultaneously capture the antigen landscape, paired TCR repertoire, and gene expression of T cell phenotypes in a high-throughput manner. Many existing methods require re-expression of receptors or ligands on heterologous cells, and thus cannot be applied directly to human clinical samples. Similar challenges apply to study BCR-antigen interactions, with the added challenge of addressing known intracellular antigen epitopes recognized by antibodies.

Here we developed a three-in-one platform—to display, deliver, record in one virion— to systematically decode ligand-receptor interactions. We engineer lentiviruses at multiple levels including (1) displaying user-defined proteins or epitopes on the viral surface; (2) engineering a fusogen to achieve receptor-specific cell entry of cognate ligand displayed viruses; (3) delivering fluorescent proteins to track engineered viruses; and (4) modifying viral RNA to record ligand information by sequencing. We termed this technology ENTER (l**e**ntiviral-mediated cell e**nt**ry by **e**ngineered ligand-**r**eceptor interaction), which can systematically deorphanize pairs of interactions including TCR-pMHC, antibody-antigen, costimulatory ligand-receptors, and intracellular antigen-BCR. We further combined ENTER with droplet-based single-cell genomics profiling (ENTER-seq) to measure antigen specificity, TCR repertoire, gene expression and surface protein landscape in individual human primary T cells.

## RESULTS

### A viral display platform to capture ligand-receptor interactions

Viruses are a natural display and delivery system. First, they infect specific cell types based on viral envelope recognition of host receptors, allowing for programmable ligand-protein display (Yang et al., 2006). Second, they assemble thousands of copies of viral proteins into one viral particle, permitting efficient protein cargo delivery (Briggs et al., 2004; Kaczmarczyk et al., 2011). Third, viruses such as retroviruses carry single-stranded viral RNA, enabling a record of genetic information (Moore and Hu, 2009). Last, their viral RNA can be reverse transcribed and integrated into the host genome, allowing for stable gene delivery (Verma and Somia, 1997) (**Figure 1A**). Here we have engineered lentivirus at multiple levels including (i) ligand proteins displayed on the viral surface, (ii) host receptor-targeted viral entry by displayed ligand and modified fusogen, (iii) fluorescent protein delivery via fusion of viral structural protein, and (iv) tagged viral RNA for single cell sequencing (**Figure 1A**).

**Figure 1.**
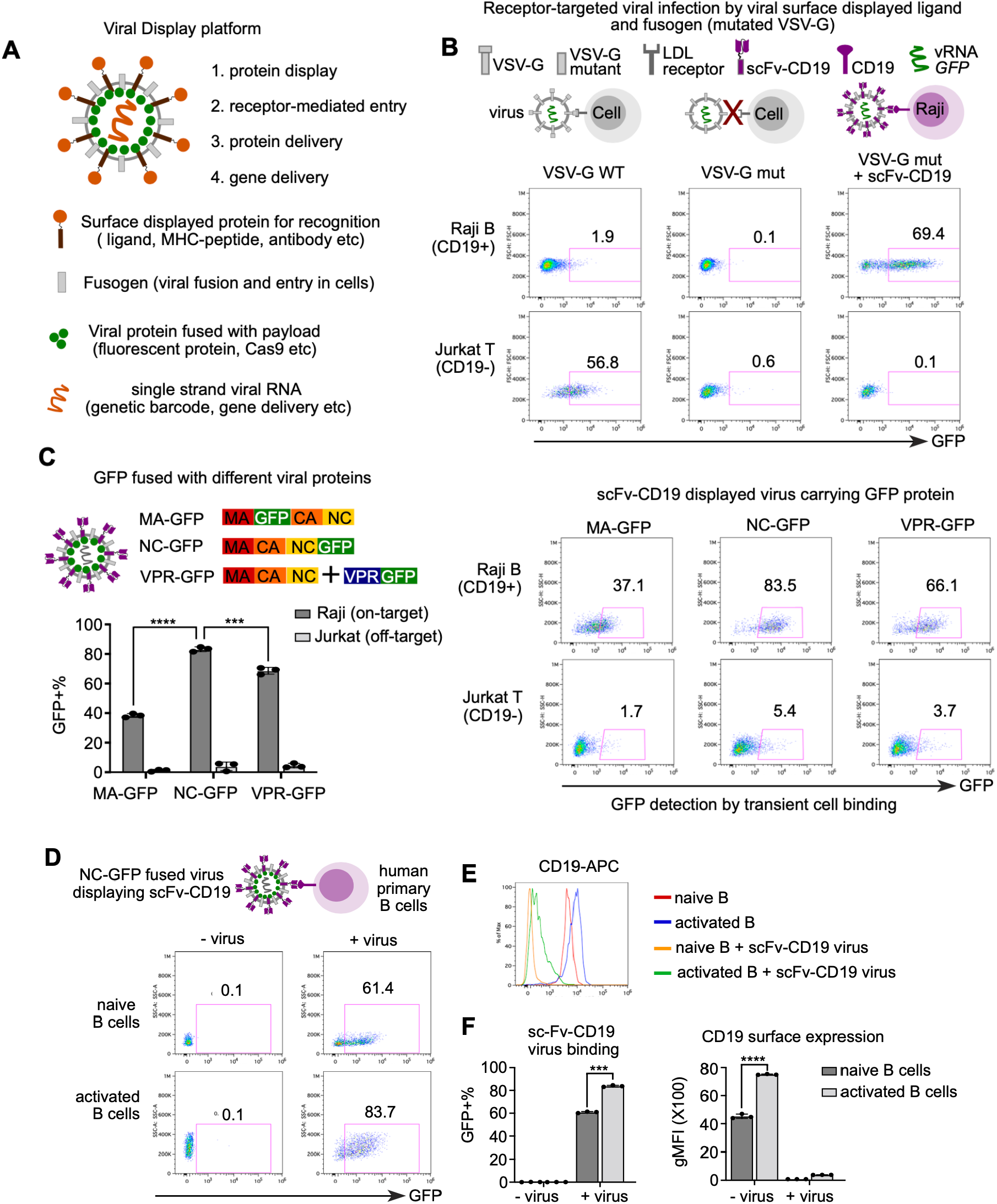
A platform to display ligand proteins and fusogen on viral surface, deliver fluorescent proteins, and record ligand-receptor interaction by cell entry. A. Schematic view of all-in-one viral platform. The lentiviruses are engineered in diverse components including: (1) user-defined ligand proteins displayed on viral surface; (2) modified fusogen with intact fusion ability and defective binding to natural receptors; (3) cargo proteins fused with viral structure protein; and (4) barcoded viral RNA for tracing and gene delivery. B. Schematic view of experimental set up and flow cytometry analysis of GFP expression after 3 days of viral infection. Raji and Jurkat cells are infected by three groups of lentiviruses encoding *GFP* in the viral RNA: (1) viruses with wild-type VSV-G (left); (2) viruses with receptor-binding mutated VSV-G (middle); (3) viruses with VSV-G mutant and anti-CD19 single-chain antibody variable fragment (scFv). C. Schematic view (left top) of experimental set up and flow cytometry analysis (right) of GFP signal after transient viral incubation. GFP protein are fused with matrix protein (MA-GFP) or Nucleocapsid protein (NC-GFP), or viral protein R (VPR-GFP). scFv-CD19 displayed viruses carrying GFP protein fused with different viral proteins were incubated with Raji (CD19+) or Jurkat (CD19-) cells for 2 hours and then subjected to flow cytometry. Bar plot (left bottom) showing the percentage of GFP+ cells upon incubation of viruses with different GFP fusion viral proteins. D. Schematic view of experimental set up and flow cytometry analysis of GFP signal in primary human B cells with or without viral incubation. Naïve and activated human primary B cells were incubated with NC-GFP fused and scFv-CD19 displayed viruses for 2 hours and then subject to flow cytometry analysis. B cells were gated on live CD20+ cells. E. Histogram analysis of surface CD19 expression of groups from Figure 1D. F. Bar plots showing scFv-CD19 virus binding and CD19 surface expression in naïve and activated human B cells *P*-values in Figure 1C and 1F are calculated by unpaired t-test. **** P<0.0001 *** P<0.001

To achieve specific ligand-receptor interaction between lentiviruses and host cells, we reason that a viral envelope protein with disrupted native receptor binding while maintaining intact fusion ability (termed as fusogen) is essential to cooperate with user-defined ligand-proteins displayed on viral surface. We envision that such cooperation of two separate modules (ligand-protein + fusogen) allows the interaction between viral-displayed ligand and host receptor to further facilitate viral fusion to host cells by fusogen (**Figure 1A**). We focused on vesicular stomatitis virus G protein (VSV-G), a viral envelope protein that has been extensively used to pseudotype lentiviruses (Burns et al., 1993). VSV-G pseudotyped viruses have a broad tropism since VSV-G can recognize and interact with Low Density Lipoprotein Receptor (LDLR), which is expressed in many cell types (Finkelshtein et al., 2013). After we infected the Jurkat T cells and Raji B cells with the VSV-G pseudotyped lentiviruses carrying *GFP* transgene, we found a robust GFP expression in these cells, although with different transduction efficiency which is potentially due to variable expression of LDLR on diverse cell types (**Figure 1B**).

We then engineered a VSV-G mutant that harbors two point-mutations (K47Q, R354A) to prevent its recognition and interaction with LDLR on host cells (Nikolic et al., 2018). As expected, we saw minimal GFP expression in Raji (0.1%) and Jurkat (0.6%) cells using mutant VSV-G pseudotyped viruses (**Figure 1B**), suggesting that the viral recognition of specific receptors on host cells is the first essential step for viral entry and integration. To test if the VSV-G mutant is a good fusogen candidate to cooperate with user-defined ligands on the viral surface for viral infection in host cells expressing paired receptors. we co-expressed a well-established CD19-CAR (chimeric antigen receptor) that contains an anti-CD19 single-chain antibody variable fragment (sc-Fv), the VSV-G mutant, the GFP transgene in a viral transfer vector, and packaging component for lentiviruses in HEK 293T cells and collected viruses from supernatant. As expected, such viruses specifically infected Raji B cells with high expression levels of CD19 but not CD19-negative Jurkat T cells (**Figure 1B**). Thus, our viral display platform reprograms the viral fusion from cell entry that is dependent on the native VSV-G/LDLR interaction to cell entry that is dependent on the interaction between user-defined ligand and paired host cell receptors.

Next, to capture ligand-receptor interaction while avoiding the viral integration and integration-induced mutagenesis of the host genome (Ranzani et al., 2013), we utilized a viral integrase mutant (D64V) that cannot integrate the host genome (Certo et al., 2011). We also fused the GFP protein with viral structural proteins to track ligand displayed viruses. We measured the transient viral entry into host cells expressing paired receptors using ligand-displayed viruses carrying GFP protein instead of viral integration to express GFP. To identify the viral protein that serves as an optimal fusion partner for GFP, we tested three viral proteins including matrix protein (MA), nucleocapsid protein (NC), and HIV accessory protein named viral protein R (VPR) (**Figure 1C**). During viral assembly and synthesis, MA and NC are processed from Gag precursor protein which can be assembled in *cis* as 3000 copies of MA or NC per viral particle (De Guzman et al., 1998; Kutluay et al., 2014). VPR can be incorporated in *trans* into the viral particle via interaction with Gag protein as 500 copies of VPR per viral particle (Wu et al., 1995). We found that 80% of Raji cells are bound by CD19-CAR presented viruses when GFP is fused with NC, which significantly outperforms MA-GFP and VPR-GFP (**Figure 1C**). Next, to investigate if the NC-GFP viruses displaying CD19-CAR can recognize and bind to primary CD19+ B cells from human blood, we incubated these viruses with naïve or activated human primary B cells for 2 hours and detected the GFP signal on these B cells by flow cytometry. Similar to Raji B cells, 80% of activated human primary B cells were bound by NC-GFP labeled CD19-CAR displayed viruses whereas 60% of naïve B cells were GFP+ (**Figure 1D**). We reasoned that the difference of CD19 expression between naïve and activated B cells might account for the discrepancy of binding of CD19-CAR viruses. Indeed, flow cytometry results showed that the surface expression of CD19 in activated B cells is significantly higher than that of naïve B cells (**Figure 1E-F**), which is consistent with the higher binding of viruses on activated B cells. This result suggests that the binding of ligand-displayed viruses to receptor-expressed cells is quantitatively correlated with the expression level of ligand-paired receptors. Moreover, CD19 surface expression was dramatically decreased after incubation of CD19-CAR viruses, indicating that CD19-CAR viruses specifically bind to CD19 to prevent the later binding of flow cytometry antibody targeting CD19, either through CD19 antigen masking or inducing internalization of surface CD19 (**Figure 1E-F**).

To further determine if our ligand-displayed GFP fused viruses can specifically bind to target cells through other ligand-receptor interactions, we engineered viruses to display either wild-type CD40 ligand (CD40L) or mutant CD40L. The CD40L mutant contains two point-mutations (K142E, R202E), leading to decreased binding affinity towards CD40 (Pasqual et al., 2018). The flow cytometry results showed a significant decrease of GFP+ Raji cells when incubated with viruses displaying CD40L mutant compared to wild-type CD40L (**Figure S1**). These results suggest that our viral display platform can capture a highly specific ligand-receptor interaction in a transient viral binding assay that avoids viral-integration induced mutagenesis of the host genome and is applicable to multiple categories of receptor-ligand interactions.

### ENTER with pMHC displaying viruses maps TCR specificity

To investigate if our viral display platform can capture the interaction between pMHC and TCR, we engineered viruses to display a single chain of MHC fused with beta 2 microglobulin (B2M) and covalently linked peptide (**Figure 2A**). To prevent interference of the endogenous human leukocyte antigen (HLA, the human MHC locus) from HEK 293T cells when producing HLA-peptide displayed viruses, we made a stable HLA knockout (KO) HEK 293T cell line by knocking out all potential HLA class I alleles (HLA-A/B/C) by CRISPR-Cas9 (**Figure S2A**). The surface expression of B2M is also abolished in HLA KO cells, suggesting all endogenous HLA alleles have been successfully deleted (**Figure S2B**). We overexpressed the single chain of HLA-A*0201 (HLA-A2) fused with B2M and peptide in HLA KO cells and observed a high level of surface expression of HLA-A2 and B2M (**Figure S2**).

**Figure 2.**
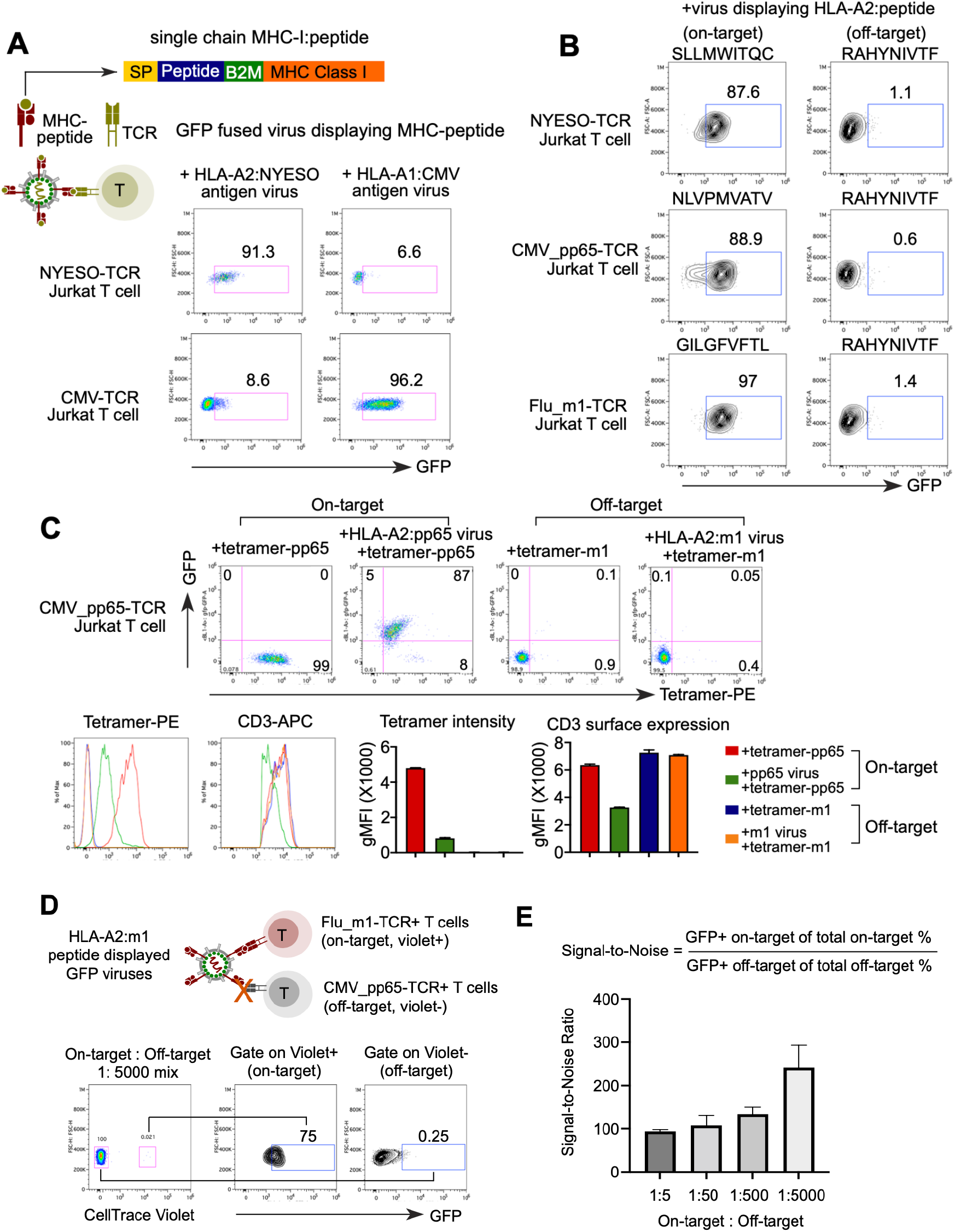
ENTER decodes interaction between pMHC with TCR. A. Schematic view of pMHC displayed viruses and flow cytometry analysis of GFP signal in specific TCR expressing Jurkat T cells upon incubation of pMHC displayed viruses. GFP fused viruses displaying either a 9-mer peptide (NYESO_157-165_) presented by HLA-A0201 allele or an 11-mer CMV peptide (pp65_363-373_) presented by HLA-A0101 allele were incubated with T cells expressing specific TCRs (e.g., NYESO_157-165_-TCR or CMV_pp65(363-373)_-TCR) that recognize the cognate antigens. SP: signal peptide; Peptide: antigen peptide; B2M: Beta-2-Microglobulin. B. Flow cytometry analysis of GFP signal in Jurkat T cells expressing specific TCRs (e.g., NYESO_157-165_-TCR, CMV_pp65(495-503)_-TCR, or Flu_m1(58-66)_-TCR) upon incubation of viruses displaying various HLA-A2 presented peptides. RAHYNIVTF peptide displayed viruses serve as a negative control. C. Flow cytometry plots (top) of CMV_pp65(495-503)_-TCR T cells upon 2-hour incubation of HLA-A2-peptide displayed GFP viruses followed by staining of PE-tetramers. M1_(58-66)_ (flu antigen) displayed viruses and M1_(58-66)_ tetramers are negative controls. Histogram plots (bottom left) and Bar plots (bottom right) showing tetramer intensity and CD3 surface expression of CMV_pp65(495-503)_-TCR T cells. D. Schematic view of experimental set up (top) and flow cytometry analysis (bottom) of TCR-T cell mixing experiment. Flu_m1(58-66)_-TCR T cells were labeled by CellTrace Violet dye and then mixed with CMV_pp65(495-503)_-TCR T cells at different ratio. Mixed T cell population was incubated with HLA-A2:m1 displayed GFP viruses for 2h and then subjected to flow cytometry. Representative flow cytometry plot showing 1:5000 mixing of two T cell population and GFP signal of T cells gated on Violet+ and Violet-population. E. Bar plot showing the signal-to-noise ratio of ENTER in Figure 2D and Figure S2B.

Next, we engineered GFP-fused reporter viruses to display pMHCs on the surface by co-expressing the single-chain of HLA-peptide, mutant VSV-G, and viral Gag protein containing NC-GFP in HLA KO HEK 293T cells. We collected the viruses and incubated them with Jurkat T cells expressing a TCR that targets the cognate pMHC antigen. Using these modular components, we successfully generated viruses displaying a well-established cancer-testis antigen NY-ESO-1 as a 9-mer peptide (SLLMWITQC) on HLA-A2, a most prevalent HLA allele in humans (Jäger et al., 1998)(**Figure 2A**). Over 90% of cognate NY-ESO-1 TCR-expressing T cells were labeled by NY-ESO-1 antigen-bearing GFP viruses, compared to 8% of T cells specific to a known human cytomegalovirus (CMV) epitope (Lee and Meyerson, 2021). Similarly, viruses displaying a CMV antigen as an 11-mer (YSEHPTFTSQY) peptide on a different HLA allele HLA-A*01:01, specifically entered into CMV TCR-T cells rather than NY-ESO-1 TCR-T cells (**Figure 2A**). We further engineered viruses displaying diverse 9-mer antigen epitopes from cancer-testis antigen, CMV pp65_495-503_ antigen, and influenza matrix protein antigen (M1_58-66_), all of which are presented on HLA-A2 allele (Gotch et al., 1987; Wills et al., 1996). The result showed over 80% of TCR-matching T cells are GFP+ after 2 hours of incubation with GFP fused viruses displaying cognate HLA-A2-peptide whereas only 1% of these T cells are labeled by negative control antigen displayed GFP viruses (**Figure 2B**). The highly specific entry of pMHC displaying viruses into TCR-matching T cells is observed with different antigen peptide lengths and distinct HLA alleles, highlighting the generality of the ENTER platform to present diverse pMHC antigens.

To further test the specificity of our pMHC displayed viruses, we first incubated the CMV pp65_495-503_ antigen targeting TCR Jurkat T cells with pp65_495-503_ displayed viruses and then stained with a widely used commercial pp65_495-503_ tetramer. For negative controls, we incubated these T cells with the influenza M1_58-66_ displayed viruses and commercial M1_58-66_ tetramer (**Figure 2C**). Flow cytometry result showed that over 90% of tetramer positive cells are GFP+, indicating a strong concordance of pp65_495-503_ tetramer staining with the binding of pp65_495-503_ displayed GFP viruses. The negative control M1_58-66_ tetramers and viruses did not label pp65_495-503_ TCR T cells (**Figure 2C**). Despite the correlation of tetramer staining with virus binding, we observed that the pp65_495-503_ tetramer intensity and CD3 surface expression are significantly decreased after co-incubation with viruses displaying pp65_495-503_.

To determine the specificity and sensitivity of our pMHC viral display platform, we mixed on-target T cells (TCR recognizing flu M1_58-66_ antigen) and off-target T cells (TCR recognizing CMV pp65_495-503_ antigen) at different ratios and then incubated with the M1_58-66_ antigen presented GFP viruses (**Figure 2D, S2B**). To distinguish these two different TCR T cell line, we labeled the flu-M1 TCR T cells with cell trace violet dye. We calculated the signal-to-noise ratio based on the frequency of on-target GFP+ cells versus that of off-target GFP+ cells. The signal-to-noise ratio was over 200-fold even when the frequency of on-target T cells is as low as 1 in 5000, demonstrating a high specificity and sensitivity of our viral display platform (>99% of specificity and up to 95% of sensitivity, **Figure 2E, S2B**).

### Decoding BCR interaction with antigen epitopes derived from intracellular proteins

The successful application of our viral display platform in capturing the interaction between TCR and pMHC encouraged us to explore the feasibility of capturing the interaction between BCR and antigens. Unlike TCR recognition of antigen peptides that are presented by MHC on the cell surface, BCR can recognize antigen epitopes that are derived from not only cell surface proteins but also intracellular proteins. To display antigen epitopes from intracellular proteins on the viral surface, we sought to engineer a transmembrane (TM) domain for optimal surface display of B cell antigen epitopes. To select candidate TM domains for viral surface display, we took advantage of the unique ability of HIV-1 viruses to incorporate host proteins on the viral surface during virus budding. Nascent HIV-1 viruses can selectively incorporate certain host TM proteins while excluding other abundant host surface proteins during viral assembly and budding (Burnie and Guzzo, 2019). We prioritized a list of highly abundant host TM proteins that are incorporated into viral surface from previous literature using mass spectrometry of viruses, immunocapture assay, and flow virometry method (Burnie et al., 2020; Cantin et al., 1996; Chertova et al., 2006; Grover et al., 2015; Jalaguier et al., 2015). This list of host TM proteins includes MHC class I and II molecules (HLA-DRA, HLA-DRB, HLA-A2), adhesion molecules (ICAM1, CD43, CD162, CD62L), and integrin family members (CD49d, LFA-1) (**Figure 3A**).

**Figure 3.**
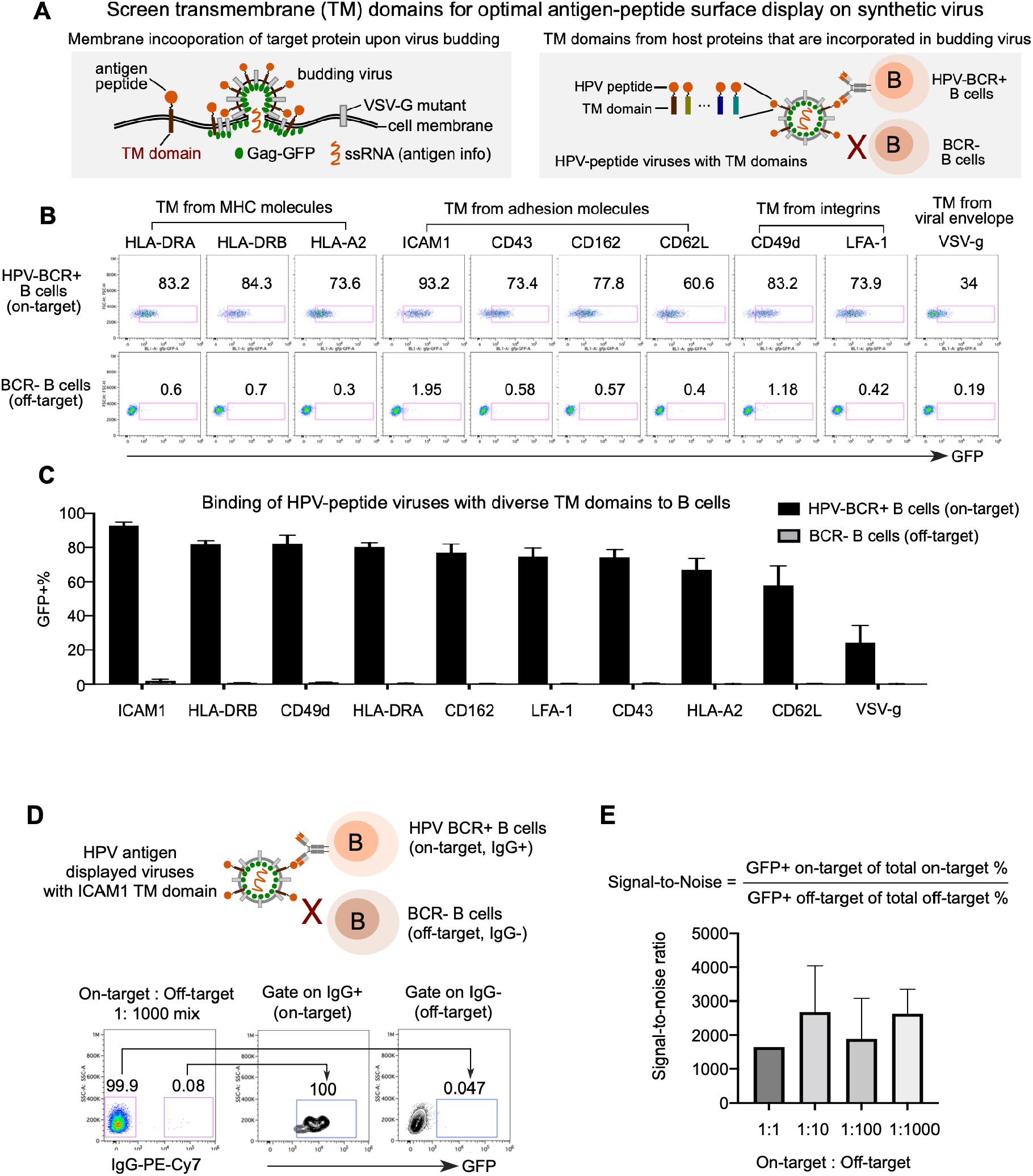
Optimization of ENTER to present intracellular antigens on viral surface and decode interaction between BCR with antigens. A. Schematic view of experimental design. During the assembly and budding of lentiviruses, certain host cell surface proteins can be incorporated into the surface of viruses. TM domains of host proteins selected from literature are fused with a B cell epitope derived from intracellular antigen HPV16 L2. These HPV-epitope displayed GFP viruses are incubated with B cells expressing BCR targeting this HPV epitope (on-target) or B cells without any BCR expression (off-target). B. Flow cytometry analysis of GFP signal in B cells incubated with GFP viruses displaying HPV epitope fused with different TM domains. B cells without BCR expression serve as a negative control. C. Bar plots showing the percentage of GFP+ B cells from Figure 3B. D. Schematic view of experimental set up (top) and flow cytometry analysis (bottom) of B cell mixing experiment. HPV-BCR+ B cells were mixed with BCR- B cells at different ratio. Mixed B cell population was incubated with GFP viruses displaying HPV epitope fused with ICAM1 TM domain and then subjected to flow cytometry. Representative flow cytometry plot showing 1:1000 mixing of two B cell population and GFP signal of B cells gated on IgG+ and IgG- population. E. Bar plot showing the signal-to-noise ratio of ENTER from Figure 3D and Figure S3.

To determine the specificity and efficiency of viral display of B cell epitopes with these diverse TM domains, we engineered viruses expressing a B cell antigen epitope derived from human papillomavirus (HPV) minor capsid antigen L2 (HPV16 L2_17-36_) and fused with TM domains from the prioritized list. Next, we generated a BCR expressing B cell line that specifically targets this HPV16 L2 B cell epitope (Wang et al., 2015). After incubating the TM domain fused and B cell epitope displayed viruses with B cells either expressing HPV-BCR (on-target) or without any BCR (off-target), we quantified the percentage of GFP+ B cells to measure the efficiency and specificity (**Figure 3B**). In addition to TM domains from host proteins, we further engineered the viruses to fuse the B cell epitope with the TM domain from the fusogen VSV-G, a viral envelope protein that can be assembled in budding viruses. The results revealed ICAM1 TM domain as the top candidate since over 90% of HPV antigen-specific BCR+ B cells are GFP+ (**Figure 3B-C**). This is consistent with previous reports showing ICAM1 is selectively acquired in budding viruses through interaction with viral matrix protein (Jalaguier et al., 2015).

To further determine the specificity and sensitivity of our viruses displaying B cell epitope fused with ICAM1 TM domain, we mixed on-target B cells (with BCR recognizing HPV L2 antigen) and off-target B cells (without BCR expression) at different ratios and then incubated with the TM-domain optimized HPV L2 antigen epitope presented viruses (**Figure S3A**). To distinguish these B cells with or without BCR, we stained these cells with IgG to specifically label B cells expressing IgG BCR. We calculated the signal-to-noise ratio based on the frequency of on-target GFP+ cells versus that of off-target GFP+ cells. The signal-to-noise ratio was over 2000-fold even when the frequency of on-target B cells is as low as 1 in 1000, indicating a profound specificity and sensitivity of our TM domain optimized viruses to display B cell antigen epitopes (>99.9% of specificity and >95% of sensitivity, **Figure S3B**).

### ENTER-seq captures pMHC specificity, TCR repertoire and gene expression profile at single-cell resolution

Given the unique feature of viral RNA that carries genetic information to encode surface-displayed ligand proteins, we combined our viral display platform with droplet-based single-cell RNA-seq to develop ENTER-seq, a technology to capture the ligand-receptor interaction and molecular blueprints at the single-cell resolution. Specifically, we applied ENTER-seq to simultaneously map pMHC antigen specificity, TCR repertoire and gene expression of T cells at the single-cell level using our pMHC-displayed viruses. The workflow of ENTER-seq includes (1) generation of a pooled pMHC-displayed GFP viruses, (2) incubation of these viruses with human T cells, (3) sorting virus-labeled GFP+ cells for droplet-based single cell profiling (e.g., 5 prime single-cell RNA-seq/V(D)J-seq by 10x Genomics), (4) generation and sequencing of three single-cell libraries including gene expression, V(D)J TCR repertoire, and antigen-peptide sequence (**Figure 4A**).

**Figure 4.**
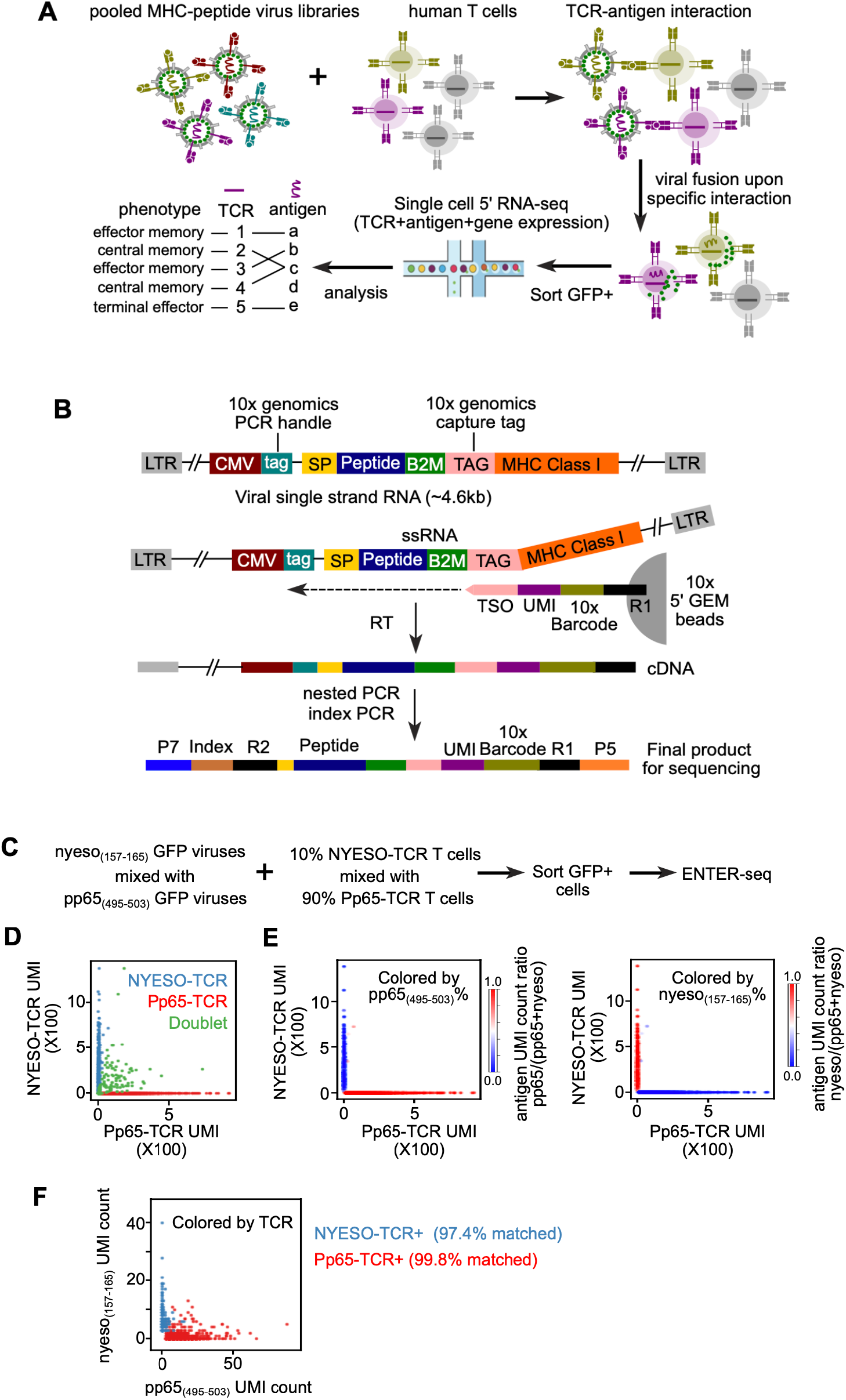
ENTER-seq for massively parallel measurement of antigen peptide sequence, TCR sequence, and transcriptome. A. Schematic view of ENTER-seq workflow. A library of pooled viruses displaying individual pMHC was incubated with T cells for 2 hours. GFP+ T cells were sorted for droplet-based single cell genomics profiling (e.g. 10x genomics 5’ kit for gene expression and V(D)J immune profiling). B. Viral RNA engineering strategy for droplet-based single cell capture. 10x genomics capture tag is inserted in the linker region between B2M and HLA-A2. 10x genomics PCR handle is inserted after CMV promoter. CMV: CMV promoter; SP: signal peptide sequence; Peptide: antigen peptide; B2M: Beta-2-Microglobulin; MHC Class I: HLA- A0201 allele; LTR: long terminal repeat; TSO: template switching oligo sequence. C. Schematic view of T cell mixing experiment for ENTER-seq. D. Number of Pp65_(495-503)_-TCR T cells UMI counts (x-axis) and NYESO_157-165_-TCR UMI counts (y-axis) associated with each cell barcode (dot). The colors are assigned as NYESO_157-165_-TCR+ T cells (light blue), Pp65_(495-503)_-TCR+ T cells (red), and doublets (green, with both NYESO_157-165_-TCR and Pp65_(495-503)_-TCR). E. Scatter plots of TCR UMI counts after doublet removal, colored by enrichment ratio of pp65_(495-503)_-antigen UMI count among total UMI counts (left), and enrichment ratio of nyeso_157-165_-antigen UMI count among total UMI counts (right). F. Number of pp65_(495-503)_-antigen UMI counts (x-axis) and nyeso_157-165_-antigen UMI counts (y-axis) associated with each cell barcode (dot) after doublet removal. The colors are assigned as NYESO_157-165_-TCR + T cells (light blue) and Pp65_(495-503)_-TCR + T cells (red).

The single-chain pMHC information is stored in viral single-strand (ss) RNAs that are packaged into viral particles. The viral ssRNA is approximately 4.6 kb, making it difficult to reverse transcribe (RT) into full-length cDNA in droplets. To efficiently capture the MHC-peptide information on viral RNA during the RT step in each droplet, we inserted a capture tag in the linker region between B2M and MHC, and another PCR handle next to the CMV promoter (**Figure 4B**). This capture tag allows capture by commercially available 5’ GEM beads through hybridizing with the Template Switch Oligo (TSO) sequence conjugated on the beads. The PCR handle permits convenient amplification of targeted peptide sequence without spiking in additional primers during the cDNA amplification step (**Figure 4B**). Further nested PCR and index PCR allow targeted enrichment of the antigen peptide sequence to generate the final antigen library for deep sequencing. The insertion of the capture tag and PCR handle does not affect the display of pMHC on viruses and specific interaction with TCR-expressing T cells (**Figure S4A-B**).

To benchmark ENTER-seq for single-cell profiling of antigen specificity and TCR repertoire, we performed ENTER-seq on mixed TCR-expressing T cells with a pooled pMHC displaying viruses. To mimic a real-life T cell population, we mixed 10% of T cells with TCR recognizing NYESO antigen and 90% of T cells with TCR recognizing CMV pp65 antigen, and then incubated with pooled viruses displaying NYESO antigen or pp65 antigen (**Figure 4C**). Analysis of the unique TCR sequence after filtering out doublets confirmed the mixing ratio of T cells at 9.4% (NYESO-TCR) vs 90.6% (Pp65-TCR), which is similar to the input mixing ratio (10% vs 90%) (**Figure 4D).** The quality control metrics for single-cell transcriptome from ENTER-seq are comparable to other published single cell RNA-seq data (Zheng et al., 2017)(**Figure S4C**). We further recovered 4198 T cells with reliable antigen peptide information and TCR sequence after filtering the unique molecular identifier (UMI) count of TCRs and antigen peptides. We calculated the ratio of the UMI from the dominant antigen peptide among total peptides and observed a high concordance of antigen peptides to their paired TCR (**Figure 4E**). After matching TCR sequences to antigen peptides at the single-cell level, the result showed that 99.8% of HLA-pp65+ cells and 97.4% of HLA-nyeso+ cells are matched with their corresponding TCR sequences respectively (**Figure 4F**). Thus, ENTER-seq can sensitively and robustly capture the interaction of TCR repertoire and cognate HLA antigen peptide at the single-cell resolution.

### Optimized ENTER-seq detects rare antigen-specific primary human T cells

To determine if our ENTER-seq can be applied to rare antigen-specific primary T cells isolated from human blood, we first validated the sensitivity of our system using GFP viruses displaying CMV-pp65 antigen epitope presented on HLA-A2 allele, and primary T cells from HLA-A2+ patients with a history of CMV infection. We incubated these primary T cells with pp65_495-503_ antigen-displayed viruses and then stained them with a widely used CMV pp65_495-503_ tetramer which serves as a positive control. The tetramer staining analysis showed that only 1% of the T cells are pp65_495-503_-specific (**Figure S4D).** 83% of the pp65_495-503_ tetramer-positive T cells are labeled by GFP viruses. To further increase the sensitivity of detection by flow cytometry, we replaced the GFP with mNeon, a monomeric green fluorescence protein that is substantially brighter than GFP (**Figure S4D**). Indeed, 98% of pp65_495-503_ tetramer-positive T cells are recovered by mNeon viruses displaying pp65_495-503_ epitope, which is significantly higher than GFP viruses (**Figure S4D-E**).

To enrich and expand the rare antigen-specific T cells for ENTER-seq, we first cultured human peripheral mononuclear cells (PBMCs) from CMV seropositive donors with CMV antigen peptides for 10 days. The peptides are processed and presented by autologous antigen-presenting cells which will stimulate CMV antigen-specific T cells for later expansion (**Figure S4F**). To further test the specificity of our viruses on peptide-enriched T cells, we expanded T cells using peptide pp65_495-503_ and then incubated these T cells with pp65_495-503_ antigen presented mNeon viruses and then stained with pp65_495-503_ tetramer. Flow cytometry analysis showed that ∼99% of tetramer+ T cells are labeled by viruses (**Figure S4G**), demonstrating a high specificity and sensitivity of our mNeon viruses to detect peptide enriched antigen-specific T cells.

### ENTER-seq of CMV patient T cells uncover donor-specific immunogenic viral epitopes

We selected 12 CMV antigen epitopes that have been shown to present on the HLA-A2 allele, and expanded T cells from 4 different CMV seropositive donors through 12 pooled CMV epitope peptides (Lehmann et al., 2020; Lübke et al., 2020; Solache et al., 1999) (Figure S4F). We prepared 12 CMV epitope presented viruses and stained the expanded T cells with the pooled mNeon viruses displaying these CMV antigen epitopes. We observed enrichment of CMV antigen-specific T cells in 2 out of 4 donors (19.4% in donor #1803, 8.78% in donor #2042) (**Figure S4H**).

Next, we performed ENTER-seq on the expanded T cells from the blood of these two donors, and labeled each donor sample with a unique hashtag antibody (**Figure 5A**). To further interrogate the phenotype of antigen-specific T cells, we combined ENTER-seq with CITE-seq through staining cells with DNA barcoded antibodies targeting cell surface proteins CD45RA, CD45RO, and IL7R (**Figure 5A**). After sorting GFP+ (CMV antigen-specific T cells) and GFP-(bystander T cells) CD8+ T cells followed by droplet-based single cell capture, we generated libraries to profile gene expression programs, CMV antigen peptides, TCR repertoires, and surface proteins (including CITE-seq proteins and hashtag proteins) in individual cells. The result showed that cells that bound with CMV antigen displayed viruses (ENTER+) are phenotypically different from cells without virus binding (ENTER-) (**Figure 5B**). Compared to ENTER-cells, ENTER+ cells were potentially protective T cells based on high expression of effector molecules such as IFNG, TNF, and cytotoxic molecules including granzymes and perforin (**Figure 5B, Figure S5C**). Additionally, the CITE-seq data showed that ENTER+ cells are mainly effector memory T (TEM) cells (CD45RO+CD45RA-) while ENTER-cells are a mixture of naïve and central memory T (TCM) cells (**Figure 5D-E**). The single cell RNA-seq data clustered all T cells into 10 clusters including: (1) naïve T cells: CD45RA+CCR7+; (2) TCM: CD45RA-CCR7+; (3): terminally differentiated effector T cells (TEMRA): CD45RA+CCR7-; (4) mucosal-associated invariant T cells (MAIT): CD45RA-CD161+CXCR6+; (5) proliferating T cells: CD45RA+KI67+; (6) IL4+ TEM: CD45RO+IL4+; (7) KLRC2+ TEM; CD45RO+KLRC2+; (8) CST7+ TEM (CD45RO+ CST7+; (9): HSP+ TEM : CD45RO+ heat shock protein (e.g. HSPA1A)+; (10): proliferating TEM: CD45RO+KI67+ (**Figure 5C**, **Figure S5A-E**). Comparison of subset frequency between donors showed that ENTER-bystander T cells are relatively similar between two donors while ENTER+ CMV antigen-specific T cells are phenotypically different between two donors, suggesting that two donors may have different immune responses to CMV antigens.

**Figure 5.**
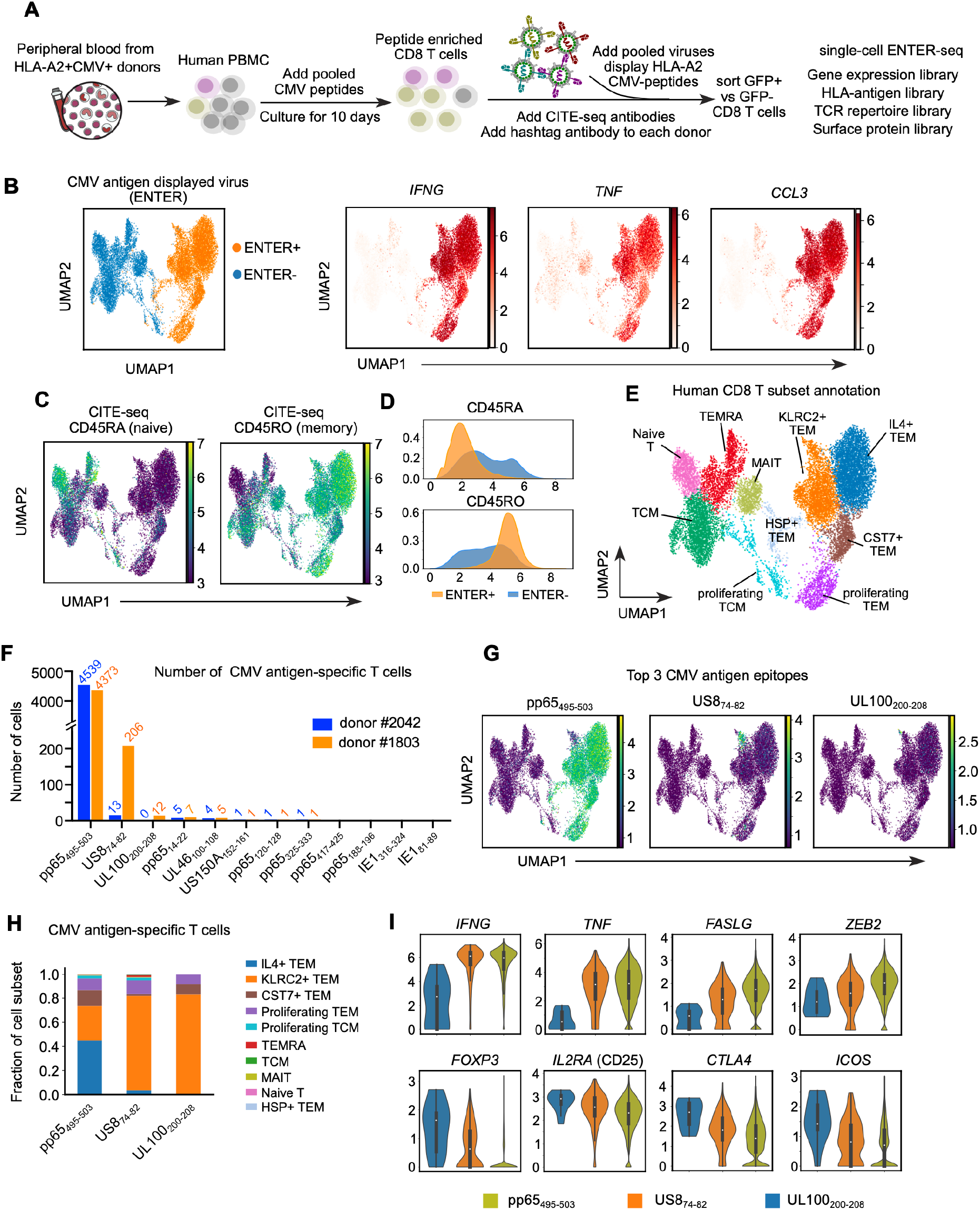
ENTER-seq of primary T cells from CMV seropositive patients. A. Schematic view of CMV antigen peptide induced T cell expansion and ENTER-seq workflow. B. UMAP plot showing cells with (ENTER+, colored in yellow) or without (ENTER-, colored in blue) CMV antigen displayed viruses binding (left). UMAP plots showing the gene expression of effector cytokines IFNG, TNF and chemokines CCL3 (right). C. UMAP plots showing CITE-seq of surface protein expression of CD45RA (naïve marker) and CD45RO (memory marker). D. Density plots showing the distribution of expression of CD45RA and CD45RO proteins in CMV antigen-specific T cells (ENTER+, colored in yellow) and bystander T cells (ENTER-, colored in blue). E. UMAP plot showing 10 clusters of human CD8+ T cell subsets labeled in different colors. F. Bar plot showing the number of CMV antigen-specific T cells that recognize specific CMV antigen epitope in donor #1803 (labeled in yellow) and donor #2042 (labeled in blue). G. UMAP plots showing the amount of CMV antigen epitopes per cell for the top 3 CMV antigen epitopes identified from F. H. Fraction of T cell subsets out of total antigen-specific T cells, separated by the top 3 CMV antigen epitopes and colored by 10 clusters of subsets from 5E. I. Violin plots showing expression of effector genes (top) or Treg genes (bottom) in CMV antigen-specific T cells, colored by top 3 CMV antigen epitope.

To determine if there is a donor-specific immune response to CMV antigens, we first measured the number of T cells that recognize specific CMV antigen epitopes in each donor. Pp65_495-503_-specific T cells are the most dominant antigen-specific T cells in both donors, suggesting pp65_495-503_ is the most common and immunogenic CMV antigen (**Figure 5F**). This is consistent with previous reports showing a high frequency of CMV pp65_495-503_ specific T cells across many donors (Elkington et al., 2003; Gillespie et al., 2000; Wills et al., 1996). We also observed a higher frequency of US8_74-82_- and UL100_200-208_-specific T cells in donor #1803 compared to donor #2042, indicating donor-specific viral epitope immunogenicity. Interestingly, when projecting the top 3 antigen peptides to gene expression UMAP plot, we observed 3 distinct clusters, suggesting different epitopes may drive unique gene CD8+ T cell fates and expression programs (**Figure 5G, 5H**). pp65_495-503_ -specific T cells span a wide range of T effector memory (TEM) fates while US8_74-82_- and UL100_200-208_-specific T cells are mostly restricted to KLRC2+ TEM fates. Moreover, pp65_495-503_ -specific T cells have high expression of effector cytokines (e.g. *IFNG*) and transcription factors essential for effector T cells (e.g. *ZEB2*) (**Figure 5I**). Surprisingly, UL100_200-208_-specific T cells have high expression of *FOXP3, CD25,* and *CTLA4*, which are characteristics of regulatory T (Treg) cells (Billerbeck et al., 2007; Churlaud et al., 2015; Fontenot et al., 2003; Wing et al., 2008), indicating that UL100_200-208_-specific T cells may resemble CD8+ Treg cells (**Figure 5I**) (Vieyra-Lobato et al., 2018). Thus, ENTER-seq not only uncovers donor-specific viral epitopes but also reveals distinct molecular blueprints of antigen-specific T cells upon recognizing different antigen epitopes from the same virus.

### ENTER-seq reveals donor-specific clonal expansion of antigen-specific T cells

To investigate the clonal expansion of CMV antigen-specific T cells, we performed an integrative analysis of TCR repertoire, antigen epitope and gene expression at the single cell level. TCR clonotypes are defined by the identity of CDR3 nucleotide sequences (Yassai et al., 2009). CMV antigen-specific T cells (ENTER+) exhibited high clonal expansion (maximum 3856 cells per TCR clone) compared to bystander T cells (ENTER-, maximum 174 cells per TCR clone) (**Figure S5A-B**). Different TCR clones from antigen-specific T cells revealed enrichment of specific TEM subsets (e.g. IL4+ TEM in TCR clone 0), whereas expanded TCR clones from bystander T cells were mainly MAIT cells or TEMRA (**Figure S6C**). Next, we asked if there is any overlap of CMV antigen-specific TCR clones between two donors. The result showed that the two donors have unique expanded TCR clones without any overlap (**Figure 6A**), consistent with prior studies showing that antigen-specific TCR clonotypes are usually private to each individual due to the high diversity of TCR repertoire (Dupic et al., 2021; Robins et al., 2009). Thus, the shared TCR specificity could not have been predicted from TCR sequences alone but required pMHC binding data. Further integrative analysis of antigen specificity and TCR clonal expansion showed that different CMV antigen epitope-specific T cells exhibit distinct behavior of TCR clonal expansion (**Figure 6B**). The clonal expansion size of pp65_495-503_ -specific T cells was significantly larger than that of UL100_200-208_-specific T cells. Highly expanded TCR clones were associated with high expression of effector genes, e.g. comparing pp65_495-503_ -specific T cells to UL100_200-208_-specific T cells. Indeed across all clonotypes, the expression of T cell effector genes was significantly correlated with TCR clonal expansion (Pearson correlation r=0.36, p=1e-100, **Figure 6C**). In contrast, the correlation between clonal expansion with other gene signatures was relatively weak (r=0.14 for T cell exhaustion genes, r=0.09 for T cell activation genes (**Figure S6D**). The correlation between clonal expansion and the expression of effector genes was observed in both donors (r=0.43 in donor #1803, r=0.33 in donor #2042, **Figure S6E**).

**Figure 6.**
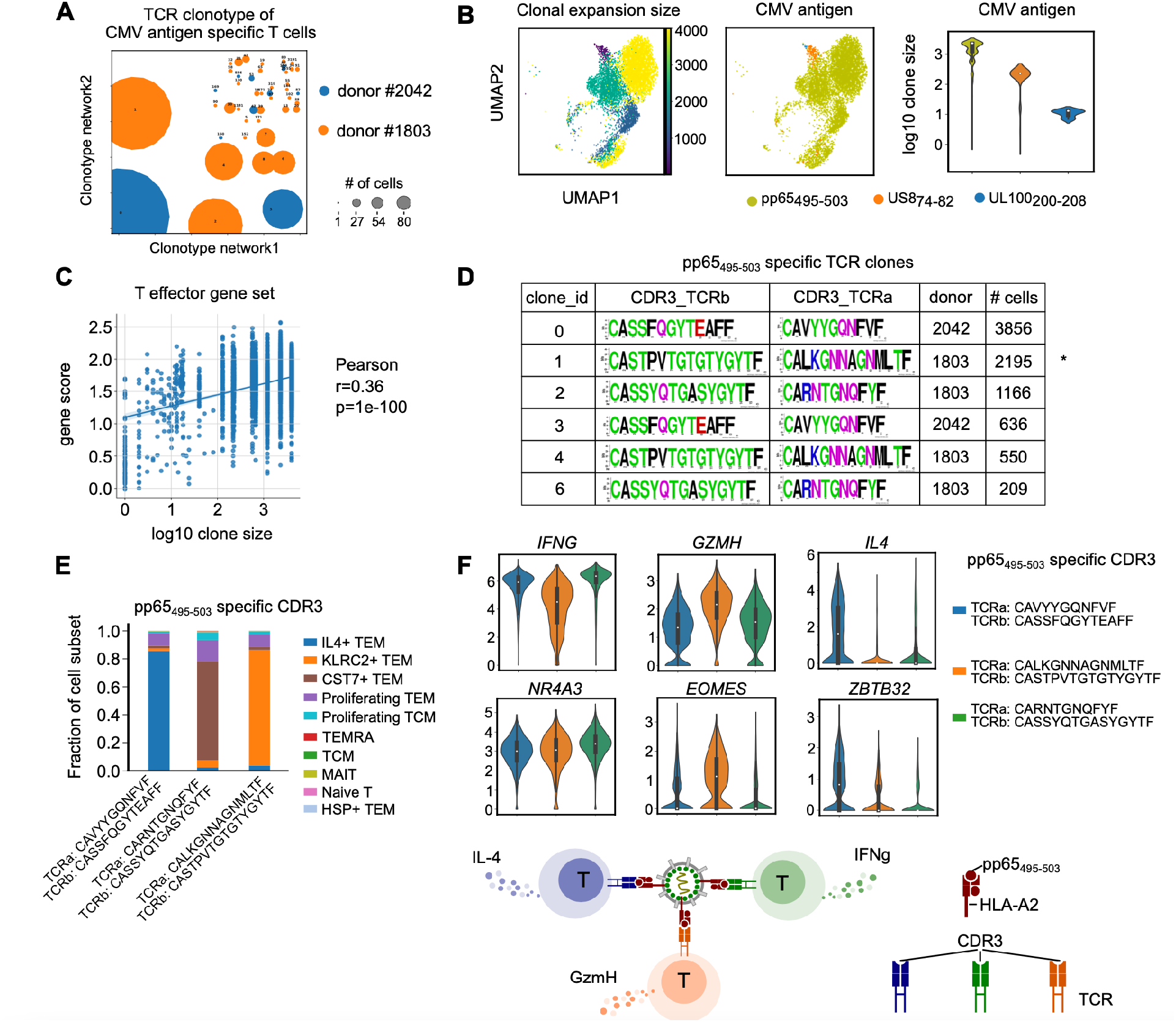
Integrative analysis of antigen specificity, TCR clonotype, and transcriptome by ENTER-seq. A. TCR clonotypes of CMV antigen-specific T cells colored by donor. Each circle represents a clonotype with identical CDR3 nucleotide sequences. The size of circle represents the number of cells in each clonotype. B. UMAP plot (left) showing the clonal expansion size of CMV antigen-specific T cells colored by the number of cells in each clonotype. UMAP plot (middle) showing the top 3 CMV antigen epitopes of antigen-specific T cells colored by antigen epitopes. Violin plot (right) showing the distribution of clone size in different CMV antigen-specific T cells colored by antigen epitopes. C. Scatter plot showing the correlation of effector gene score and the clone size in antigen-specific T cells. The fit line is added by fitting into linear regression model. The coefficient r and p value are calculated by Pearson correlation. D. A summary table of pp65_495-503_-specific TCR clones showing convergent TCR clonotypes with identical CDR3 amino acid sequences. E. Fraction of T cell subsets in pp65_495-503_-specific TCR clones, separated by CDR3 clones and colored by 10 clusters. F. Violin plots showing expression of cytokines and transcription factors in pp65_495-503_-specific TCR clones, separated and colored by CDR3 clones. A simplified model for F.

Because multiple TCR nucleotide sequences can encode the same CDR3 amino acid sequences which target the same antigen epitope, we merged the clonotypes based on identical CDR3 amino acid sequences for each CMV antigen epitope (**Figure 6D**, **Figure S6F**). For the most immunogenic CMV epitope pp65_495-503_, we identified three dominant CDR3 clonotypes. Among them, two TCR beta chain sequences (CASSFQGYTEAFF and CASSYQTGASYGYTF) are identical with published pp65_495-503_-specific TCRs in the IEDB database, further validating the specificity of our platform (**Figure 6D**) (Vita et al., 2019). Combining CDR3 clonotypes with gene expression profiles in pp65_495-503_ specific T cells, we discovered that different clonotypes exhibit distinct gene expression phenotype including distinct cytokine profiles, cytolytic enzyme, and transcription factor expression, indicating an inter-clonal phenotypic diversity that targets the same antigen epitope (**Figure 6E-F)**. Thus, ENTER-seq can functionally characterize both the TCR binding specificity and the TCR-associated cell states.

## DISCUSSION

### Three-in-one platform for ligand display, cargo delivery, and interaction recording

Here we describe the design and performance of ENTER, a versatile platform that enables three critical functions to decode cell-cell interactions. First, we engineer lentivirus to enable the display of heterologous cell surface proteins and intracellular epitopes, including pMHC complexes, antibodies, intracellular antigens. The same tethering strategy for intracellular antigens is also applicable to secreted proteins such as growth factors or cytokines. ENTER has several advantages compared to yeast or phage display platforms. The glycosylation pattern in yeast/phage display platform is different from mammalian system, which may interfere with the correct MHC presentation and recognition of paired TCRs (Wildt and Gerngross, 2005; Wolfert and Boons, 2013). Moreover, yeast or phage display requires substantial optimization to achieve proper folding, stability, and presentation of MHC (Garboczi et al., 1992). ENTER is built in human cells and enables human glycosylation and protein folding patterns, as evidenced by our ability to present multiple HLA-peptide combinations. In addition, soluble recombinant TCR is required for screening in yeast and phage display platforms, thus making it challenging to test diverse TCRs in parallel (Davis and Boyd, 2019). In contrast, ENTER allows investigators to screen primary T cell samples, opening the door to examine the vast diversity of human TCR repertoire.

Second, ENTER is a platform for RNA or protein delivery. We engineered the lentivirus such that receptor-ligand interaction drives viral fusion. We further engineered the system so that investigators may choose to integrate reverse-transcribed cDNA into the receiver cell genome or not. ENTER may have applications in gene therapy or RNA medicine as ENTER can achieve exquisite cell type sensitivity compared to existing modalities like AAV (Buchholz et al., 2015; Srivastava, 2016). We focused on non-integrative delivery of cargo protein due to several advantages. Lentiviral integration and subsequent gene expression are days-long processes and subject to multiple host cell restriction systems, which may affect the accuracy of enumerating cell numbers or states based on receptor-ligand interactions. Genome integration also has the risk of unwanted mutations, raising safety concerns for certain applications such as adoptive cell therapy (Fraietta et al., 2018; Milone and O’Doherty, 2018; Ranzani et al., 2013). In contrast, ENTER with nonintegrative cargo delivery is not confounded by cell type or state differences in integration or transgene expression, and does not permanently alter the target cell genome that complicates downstream applications (Apolonia et al., 2007). Third, ENTER has advantages over cytolytic T cell reporter assay such as T-scan because the latter cannot record the pairing of pMHC vs. TCR at single cell level (Kula et al., 2019).

### Linking ligand-receptor interaction with molecular blueprints at single-cell resolution

ENTER-seq combines the ability to decode ligand-receptor interactions with the power of single-cell genomics to resolve cell type and cell states at a massively parallel scale. ENTER-seq for pMHC is conceptually similar to a DNA-barcoded library of pMHC tetramer molecules but with several potential advantages (Bentzen and Hadrup, 2017). pMHC tetramer libraries require individual peptide synthesis and then loading into pMHC tetramers, leading to high cost, long lead time, and lower throughput compared to ENTER-seq library that can be prepared by massively parallel DNA synthesis. DNA conjugation to pMHC tetramers may suffer from unequal barcode oligonucleotide loading during the conjugation reaction, whereas ENTER-seq library leverages lentiviral biology that ensures 2 copies of barcoded viral RNA for each viral particle (Moore and Hu, 2009). Finally, ENTER-seq might be more sensitive compared to pMHC tetramers. HIV-based lentiviral particle displays 14-100 molecules of envelope protein per viral particle whereas pMHC tetramers are four linked molecules by definition (Stano et al., 2017). We further engineered the fusion of brighter mNeon into ENTER to increase its sensitivity. Together, ENTER-seq allows investigators to record ligand-receptor specificity and read out the biological consequences of this interaction, such as antigen-dependent T cell fates including naïve cell activation, effector cell expansion, memory cell formation, or T cell exhaustion. Similarly, ENTER-seq may be used to understand the molecular programs of autoantibody-producing B cells in autoimmunity.

ENTER-seq analysis of CMV patient T cells showcases the power of the platform to connect the landscape of antigen epitope, TCR repertoire, gene expression program, and surface protein phenotypes across tens of thousands of primary T cells in a single experiment. Such massively parallel profiling of diverse modalities uncovered donor-specific antigen specificity and immunogenicity, and donor-specific TCR clonal expansion in response to the same antigens. We further discovered that antigen-specific clonal expansion is associated with cytolytic effector function based on the expression of effector genes. Our ENTER-seq provides insights into a comprehensive understanding of how T cell clonality and specificity influence the molecular phenotypes and physiological function of antigen-specific T cells.

### Translational application of ENTER in immunology and beyond

We envision several translational opportunities to build on the ENTER technology. ENTER may be used to isolate and enrich tumor antigen-reactive T cells to infuse back into patients (Rasmussen et al., 2010). The nonintegrative nature of ENTER facilitates adoptive T cell therapy (Hinrichs and Rosenberg, 2014). ENTER may be furthered used in a discovery context to screen immunogenic antigen or elite TCRs for the rational design of vaccine development or cancer immunotherapy (Chandran and Klebanoff, 2019; Hinrichs and Rosenberg, 2014; Hu et al., 2018). Finally, ENTER enables cell-type-specific delivery of gene, RNA, or protein, possibly allowing immunoreceptor-specific cell transduction or gene editing (Frank and Buchholz, 2019; Hamilton et al., 2021). If ENTER may be extended to additional receptor-ligand pairs, such as G-protein coupled receptors, adhesion molecules, or protocadherins, ENTER may be used to address cell-cell connectivity beyond the immune system.

## METHODS

### Plasmid cloning and construction

Primers were ordered from IDT DNA technologies, and gene fragments were synthesized by twist biosicence and IDT. Table S1 shows the list of vector designs used in this study. All the constructs were made by gibson assembly (New England Biolabs) in general. Briefly pMD2.G (addgene#12259) was digested with EcoRI to remove wild-type VSV-g gene fragment. It assembled with mutated VSV-g (K37Q, R354Q) introduced by PCR primers to generate the VSV-g double mutant. psPax2 (addgene#12260) was digested with BsiWI and SphI to fuse eGFP after MA. To generate packaging vector with NC-eGFP/NC-mNeon fusion, psPAX2-D64V-NC-MS2 (addgene#122944) was digested with SphI and BspEI sequentially. Then part of gag and eGFP or mNeon were assembled together with backbone. GFP-VPR is obtained from Addgene (#83374)

To generate HPV16 L2 antigen specific BCR, light chain and heavy chain were amplified separately from vector JWW-1 (addgene#66748) and connected by a 2A peptide. Then it was inserted into a piggybac vector after CMV promoter (PB-CMV), after which, a PDGFR transmembrane (TM) domain and 2A-mCherry were added to express this antibody on cell surface. Single chain format of NYESO TCR (Clone 1G4, alpha and beta chain in tandem linked by a 2A peptide) was synthesized and inserted into a lentiviral vector with hygromycin resistance. TCR5, which binds to a p5 peptide from CMV virus was amplified from alpha (addgene#164999) and beta chain (addgene#165000) and made into single chain form as with NYESO TCR above.

For displaying antigen and HLA peptide complex on viral surface, we first generated a cloning lentivirus vector with a strong CMV promoter, multiple cloning sites and the WPRE element to enhance the expression. CD19-CAR vector was generated by inserting a scFv CD19 (kindly provided by Mackall lab) with a CD8 stalking linker and TM into the lentiviral plasmid followed by 2A-puromycin and 2A-eGFP. We replaced scFv CD19 and TMs to generate other antigen candidates including HPV-L2 antigen, CD40L and CD40L mutant. For TM domain screening, we swapped TM with 10 alternatives in the HPVL2 antigen viral vector (Table S2).

To display MHC-peptide complex, we built a single chain vector, which consists of a signal peptide, antigen peptide, a G4S linker, B2M, a second G4S linker and HLA allele in tandem. DNA that encodes human growth hormone signal peptide to beta2 microglobulin was synthesized and inserted into lentiviral vector together with HLA allele. Here HLA allele A0201 was amplified from addgene vector #119052, and allele A0101 was from addgene #165009. Two cysteine mutations were introduced to stabilize the peptide binding by a bisulfide bond between Y84C of HLA allele and G2C that lies in the G4S linker after peptide. To adapt it to 10xgenomics sequencing planform, we further inserted a 10xTSO sequence (Table S4) in the linker between B2M and HLA encoding amino acids SHIRN and a 10xPCR handle in 5’UTR after CMV promoter (Table S4). A cloning vector was built by replace antigen peptide with 2 esp3I sites, where various HLA peptides (Table S3) can be inserted conveniently.

### Transfection and lentiviral production

To generate regular lentivirus for cell line infection and production, per 6-well, HEK293T were transfected with a viral expression vector (2ug), pMD2.G (VSV-G wild type) (1ug), and psPax2 (2ug) with lipofectamine 3000. The media was changed one next day, and viral supernatant were collected twice at 48 hr and 72hr respectively. The virus was concentrated with 4x Lenti-X according to manufacturer’s protocol, and stored at 20x concentrated in −80c. For making specific receptor targeting and integrating virus, VSV-G mutant was used instead. To producing antigen displaying virus that can be detected with fluorescence without integration, VSV-G mutant and fluorescent protein fused version (NC-eGFP or NC-mNeon) of psPAX2-D64V (D64V mutation on integrase) vectors were mixed with antigen expressing vector (either antigen domain or HLA-peptide complex) according to above ratio to transfect the HEK293T cells. Virus were collected, concentrated to 40x, and stored in -80c.

### Cell culture and cell line production

Raji, Ramos, and Jurkat related cell lines are cultured in RPMI supplemented with 10% FBS (Invitrogen) and 1X pen/strep. HEK 293T cells are maintained in DMEM supplemented with 10% FBS and 1X pen/strep. Ramos cells are obtained from Dr Lingwood’s lab. Jurkat TCR negative -76 cells and Jurkat expressing CMV-, and flu-TCR were obtained from Dr. Mark Davis’s lab. To generate stable cell lines including BCR and TCR expressing cells, Ramos or Jurkat cells were infected with viruses, and selected by sorting or using drug after 4-5 days.

### Lentiviral infection and viral incubation assay

For Figure 1B, 30uL concentrated lentiviruses are added into 250K Raji or Jurkat cells in 12 well plate. 3 days later, GFP signal is measured by flow cytometry. For figures after 1B, 200K target cells were collected in tubes and the supernatant is removed after centrifugation. The cell pellet was resuspended in 30uL concentrated GFP fused lentiviruses and incubated at 37C. After 2hr incubation, cells are stained with flow cytometry antibodies for 10 min at 4C (if needed), washed by RPMI medium for twice, and finally subjected to flow cytometry.

### Lentivirus incubation with mixed cell population

For T cell mixing experiment, Flu-TCR expressing Jurkat T cells were labeled by CellTrace Violet dye (#C34571, Thermofisher) according to manufacturer’s protocol. Violet labeled Flu-TCR T cells were mixed with CMV-TCR T cells at diverse ratios including 1:5, 1:50, 1:500, 1:5000. The mixed T cells were incubated with 40uL concentrated HLA-A2-Flu antigen displayed GFP viruses for 2 hr at 37C. T cells were stained with CD3-APC (clone HIT3a, BioLegend) antibody, washed twice, and subjected to flow cytometry. For B cell mixing experiment, HPV-BCR expressing Ramos B cells were mixed with Ramos B cells at diverse ratios including 1:1, 1:10, 1:100, 1:1000. The mixed cells were incubated with 40uL concentrated HPV-antigen-TM (ICAM1) displayed GFP viruses for 2hr at 37C. B cells were stained with IgG-PE-Cy7 antibody (clone G18-145, BD Biosciences), washed twice, and subjected to flow cytometry. The metrics are calculated below:

Sensitivity = percentage of GFP+ on-target cells among total on-target cells

Specificity = 1- (percentage of GFP+ off-target cells among total off-target cells)

Signal-to-noise ratio = (percentage of GFP+ on-target cells among total on-target cells)/(percentage of GFP+ off-target cells among total off-target cells)

### Human primary immune cell isolation and activation

For Figure 1D, buffy coats from healthy donors were obtained from Stanford Blood Center with consent forms. Peripheral blood mononuclear cells (PBMC) were isolated using Lymphoprep (Cat# 07811, STEMCELL Technologies) density-gradient centrifugation and cryopreserved and stored in -80C. B cells were purified from thawed PBMCs by negative selection using EasySep Human B Cell Enrichment Kit (Cat#19844, STEMCELL Technologies) according to the manufacturer’s protocol. Isolated B cells were cultured in IMDM medium supplemented with 10% FBS and 55 mM beta-mercaptoethanol at 1X106 cell/mL and activated by CellXVivo Human B cell expander (1:250 dilution, R&D system) and 50 ng/mL IL2 (Cat#200-02-10ug, PeproTech) for two days. For Figure S5A, LRS chambers from HLA-A2+ donors with CMV infections (CMV seropositive) were obtained from Stanford Blood Center with consent forms. PBMCs were isolated and stored as above. CD8+ T cells were purified from thawed PBMCs by negative selection using EasySep Human CD8+ T Cell Enrichment Kit (Cat#19053, STEMCELL Technologies) according to the manufacturer’s protocol.

### Peptide enrichment of antigen-specific T cells

Short 9-mer peptides encoding CMV epitopes (compatible to HLA-A2 allele, Table S3) were synthesized by Elimbio in lyophilized powders. The peptides were dissolved in DMSO in 10mg/mL. PBMC were isolated from donor blood described as above. PBMC were cultured in T cell medium (RPMI medium supplemented with 10% FBS, 1X penstrep, 100mM HEPES, 55 mM beta-mercaptoethanol). Individual peptide (10ug/mL) or pooled peptides (1ug/mL for each peptide) were added into PBMC for culturing 10 days in T cell medium. 50ng/mL IL-2 were added every two days. After peptide enrichment, PBMCs were incubated with viruses and/or PE-tetramer and then analyzed by flow cytometry.

### Flow cytometry

For Figure 1D, B cells were incubated with viruses for 2 hr and then stained with Human TruStain FcXTM (Fc Block, BioLegend), CD19-APC (clone HIB19) and CD20-V450 (clone L27) antibody in cell staining buffer (BioLegend) for 10 min at 4C. For Figure 2, Jurkat T cells were incubated with viruses for 2 hr and then stained with CD3-APC (clone HIT3a) and PE labeled tetramer loaded with peptides (NIH tetramer core) for 30 min at 4C. For Figure S5, cells were incubated with viruses for 2 hr and then stained with human Fc Block, CD3-APC, CD8-BV711 (clone SK1), tetramer-PE if needed, and viability dye for 30 min at 4C. After staining, cells were washed twice by cell straining buffer and analyzed by flow cytometry (Attune, Thermofisher). All antibodies are from BioLegend if not specified.

### ENTER-seq workflow of mixed TCR-expressed Jurkat T cells

All primers were synthesized and ordered from IDT (table S4). Jurkat cells expressing different TCRs were mixed together and stained a mixture of virus as described above. The GFP+ cells were sorted on BD Aria II afterward. Commercial 10xgenomics 5’ RNA kit was customized to read out HLA peptide, TCR, and transcriptome simultaneously per single cell. Immediately after sorting, the cells were washed once at 4C with PBS+0.4% BSA, mixed with RT (reverse transcription) mixture spiked in with customized TCRalpha RT primers at 0.1uM, and loaded to 10x chromium. The cDNA was amplified and cleaned up to generate the transcriptome according to manufacturer’s protocol.

During cDNA cleanup, the supernatant that contains shorter fragment of HLA peptide and TCR information was further mixed with SPRISelect beads to 0.9x, and cleaned up. The library that encodes HLA peptide were generated through 2 round of nested PCRs and a final round of indexing PCR. First we enriched the HLA peptide cDNA by 8 cycles of PCR (98c for 45 min, then 8x of 20sec at 98c 20 sec, 20 sec at 59c and 30sec at 72c) with 0.5uM 10xT5_5pRNA_Fw, and GH_sp_nested1_fw. After cleanup, 5ul of elution was used the second round of PCR with 0.5um nested primer and Illumina adapters P7_Tru_GH_HLA_fw and P5_T5_inFw as above. Last, take 5ul of elution to generate final library with Illumina Truseq based index primers. The above primers were designed in a way compatible with dual index, thus either customized index primer or 10x dual index primers can be used here.

To read out TCR information in Jurkat cell lines, we generated the library that covers VDJ part of TCR alpha to infer cell’s TCR identity. First, the TCR DNA were enriched by nested PCR (specifically, 98c for 45 min, then 8x of 20sec at 98c 20 sec, 20 sec at 59c and 30sec at 72c) with 0.5uM 10xT5_5pRNA_Fw and 0.5uM mix of nyeso_TRAc_set1_rev and hs_TRAc_set1_rev targeting two different TCR. Next, take 5ul of eluent to run the second round of nested PCR with Illumina adapter (a mixture of T7TRCa_nyeso_Rev and T7TRAc_hu_Rev) and P5_T5_inFw for 8 cycles, followed by a final index PCR similar to HLA libraries.

### ENTER-seq analysis of mixed TCR-expressed Jurkat T cells

The libraries are sequenced using Illumina’s Novoseq and Miseq platforms. Transcriptiome fastq files were analyzed using 10X’s cellrangers to provide single cell barcodes. The fastq files of TCR libraries were mapped to TCR alpha chain with custom python script. The UMI count of each type TCR per cells’ barcodes was calculated. To exclude doublet, we require, per gem barcode, the UMI count for the dominant TCR is at least 10 times more than those non-dominant TCR species. Next, peptide information was extracted from the HLA library reads using custom python script to correct the UMI “hopping”, and mapped to single cell barcodes obtained above to generate a count table containing cell’s barcodes, and UMI count for TCR and HLA peptide variants. Downstream analysis and plots were generated with matplotlib package in python.

### ENTER-seq workflow of primary T cells from CMV seropositive donors

Peptide stimulated donor PBMC were collected and stain with a mixture of 12 viruses displaying CMV antigen epitopes including IE1_81-89_, IE1_316-324_, US150A_152-161_, US8_74-82_, UL100_200-208_, UL46_100-108_, pp65_417-425_, pp65_325-333_, pp65_188-196_, pp65_120-128_, pp65_495-503_, and pp65_14-22_. The peptide sequences for CMV antigens were listed in Table S3. After 2 hr, cells were also stained with barcoded antibodies CD45RA, CD45RO, and IL7R (Biolegend totalseq-C, cat# 304163, cat# 304259, cat# 351356), live dead dye, CD3-APC and CD8-BV711 for 20 min on ice. Sample from each donor is also stained with unique hash-tag barcoded antibody (Biolegend totalseq-C, cat# 394661, cat# 39466310). After two washes, CD8+ CD3+ GFP+ cells were sorted to run on 10x genomics platform using 5’ RNA and VDJ kits according to manufacturer’s protocol. Here per sample, we obtained 10x gene expression library, VDJ library, and feature barcoded CITE-seq library according to manufacturer’s protocol. In addition, HLA peptide library was generated in the same way as described above. Final libraries are sequenced on either 2X75 Miseq or 2X150 Novo-seq.

### ENTER-seq analysis of primary T cells from CMV seropositive donors

The scRNA-seq reads were aligned to GRCh38 genome and quantified using cellranger count (10x genomics). The CITE-seq reads were processed using cellranger count with antibody oligo barcode as feature reference. The TCR-seq reads were mapped to VDJ compatible reference (refdata-cellranger-vdj-GRCh38-alts-ensembl-5.0.0) using cellranger vdj (10x genomics). HLA peptide reads were processed as described above.

The later analyses for single cell RNA-seq and CITE-seq were performed using SCANPY (Wolf et al., 2018). Cells with less than 200 genes detected or greater than 10% mitochondrial RNA reads were excluded from analysis. Doublet cells were removed using CITE-seq analysis of barcoded hashtag antibody labeling donor origins. For cell clustering, raw UMI counts were first normalized by total counts to correct library size and then log-normalized. Variable genes were called using scanpy.pp.highly_variable_genes() with default parameters. Variable TCR genes were removed before principal component analysis (PCA) to prevent clustering bias from variable TCR transcripts. Next, we regress out effects of total counts per cell and the percentage of mitochondrial genes, and then scale the data to unit variance. Scaled data were used as input into PCA analysis on the basis of variable genes (without TCR genes). Clusters were identified using Leiden graph-clustering method with the first 40 principal components and resolution=0.6. UMAP plots were generated using scanpy.tl.umap() and scanpy.pl.umap() with default parameters.

Initial clusters were annotated using expression of known markers including CD3E, CD4, CD8A, CD45RA, CD45RO, CCR7, GZMB, and KLRB1. All CD8+T cells are CD3E+CD8A+CD4-. Naïve T cells are CD45RA+CCR7-. Central memory T cells (TCM) are CD45RA-CCR7+. Effector memory T (TEM) cells are CD45RO+CCR7-. Terminal effector cells re-expressing CD45RA (TEMRA) are CD45RA+CCR7-GZMA+. MAIT cells are KLRB1+CXCR6+TRAV1-2+. More specific clusters in TEM were further annotated by identifying differentially expressed marker genes for each cluster including MKI67, IL4, KLRC2, CST7, and HSPA1A. The gene score was calculated using scanpy.tl.score_genes() with ctrl_size=500 and use_raw=True. The gene set of effector genes, exhaustion genes, and activation genes for T cells were selected from previous literature (Yost et al., 2019).

TCR relevant analyses were performed using Scirpy (Sturm et al., 2020). The contig annotation files generated by cellranger vdj were used as input for TCR analysis. TCR qualities were analyzed using scirpy.tl.chain_qc(). The TCR clonotypes were defined using scirpy.pp.ir_dist() and scirpy.tl.define_clonotypes() with default parameters based on CDR3 nucleotide sequence similarity. The TCR clonotypes were visualized on a network using scirpy. tl.clonotype_network() with min_cells=3. The CDR3 amino acid compositions were generated using weblogo (Crooks et al., 2004).

All other plots (e.g. violin plots, scatter plots, density plots and bar plots) were generated by Python matplotlib and seaborn.

## ACKNOWLEDGEMENT

Supported by NIH RM1-HG007735 (H.Y.C.), the Parker Institute for Cancer Immunotherapy (H.Y.C., A.T.S.), a Career Award for Medical Scientists from the Burroughs Wellcome Fund (A.T.S.), a Technology Impact Award from the Cancer Research Institute (A.T.S.), and an ASH Scholar Award from the American Society of Hematology (A.T.S.). H.Y.C. and M.M.D. are Investigators of the Howard Hughes Medical Institute. J.A.B was supported by a Stanford Graduate Fellowship and the National Science Foundation Graduate Research Fellowship under Grant No. DGE-1656518.

## AUTHOR CONTRIBUTIONS

B.Y, Q.S., and H.Y.C. conceived the project. B.Y. and Q.S. performed experiments and analyzed the data. J.A.B, K.E.Y, and K.R.P assisted the computational analysis. H.H, M.M.D, and D.L. provided essential cell lines. H.Y.C and A.T.S. guided data interpretation. B.Y., Q.S. and H.Y.C. wrote the manuscript with input from all authors.

---

**Figure S1.**
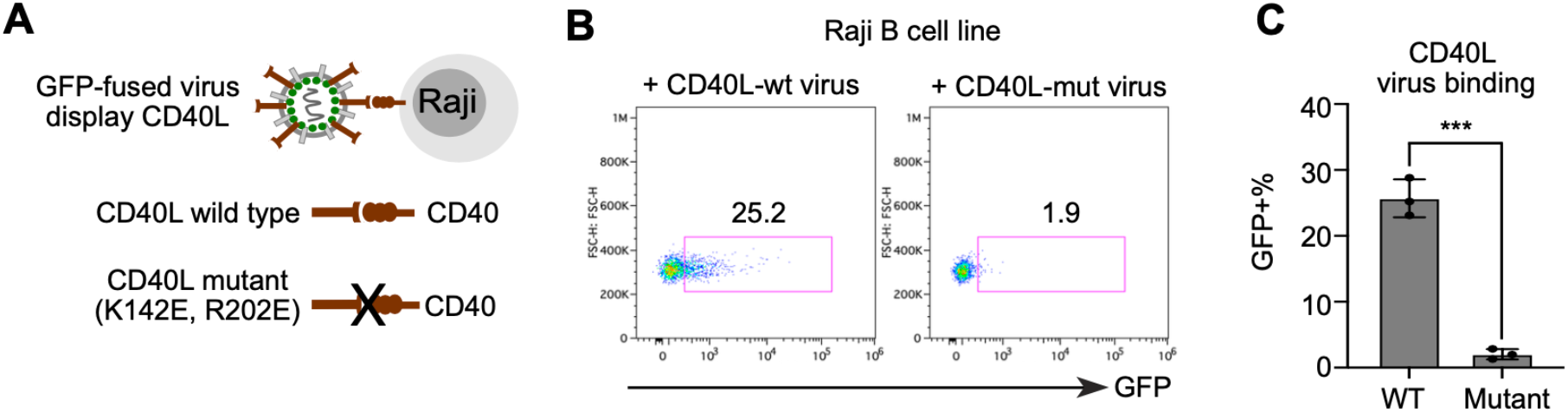
Decipher specific interaction of ligand-receptor for co-stimulatory molecules by ENTER. A. Schematic view of experimental set up. CD40 expressing Raji B cells were incubated with GFP viruses displaying either wild-type CD40 ligand (CD40L) or CD40L mutant (K142E, R202E) with decreased binding to its cognate receptor CD40. B. Flow cytometry analysis of GFP signal in Raji B cells upon incubation with either wild-type CD40L or mutant CD40L displayed GFP viruses. C. Bar plot showing the percentage of GFP+ cells from Figure S1B. P value was calculated by unpaired t-test. *** P<0.001

**Figure S2.**
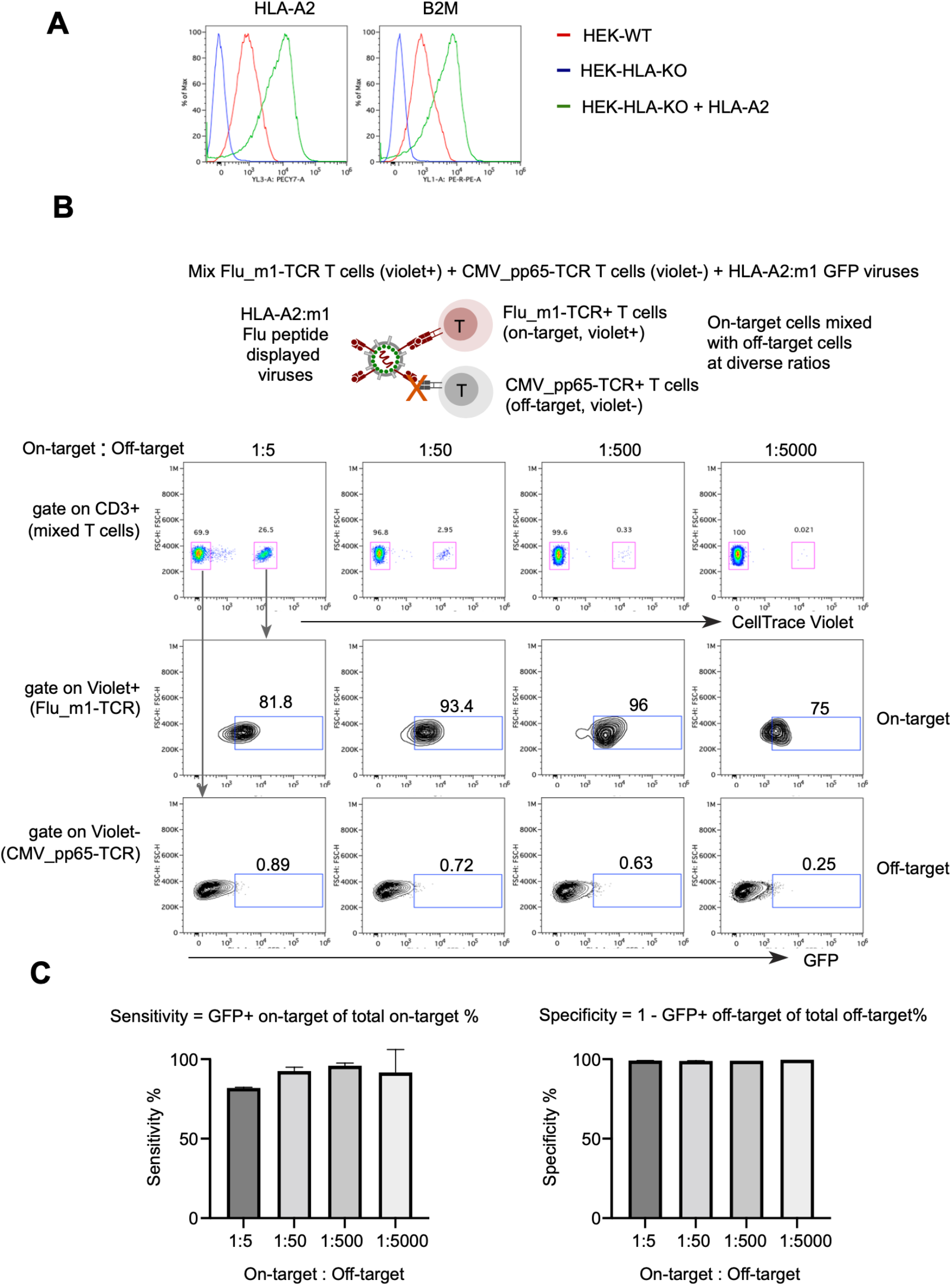
Specificity and sensitivity of ENTER in decoding interaction between TCR and pMHC. A. Flow cytometry analysis of HLA-A2 and B2M surface expression in wild-type HEK293T, HLA-KO HEK293T, and HLA-A2 reconstituted HLA-KO HEK293T cells. B. Schematic view of experimental design (top) and flow cytometry analysis (bottom) of T cell mixing experiments. Flu_m1(58-66)_-TCR T cells were labeled by CellTrace Violet dye and then mixed with CMV_pp65(495-503)_-TCR T cells at different ratio. Mixed T cell population was incubated with HLA-A2:m1 displayed GFP viruses for 2h and then subjected to flow cytometry. Representative flow cytometry plot showing mixing of two T cell population and GFP signal of T cells gated on Violet+ and Violet-population. C. Bar plots showing sensitivity (left) and specificity (right) of ENTER from Figure S2B.

**Figure S3.**
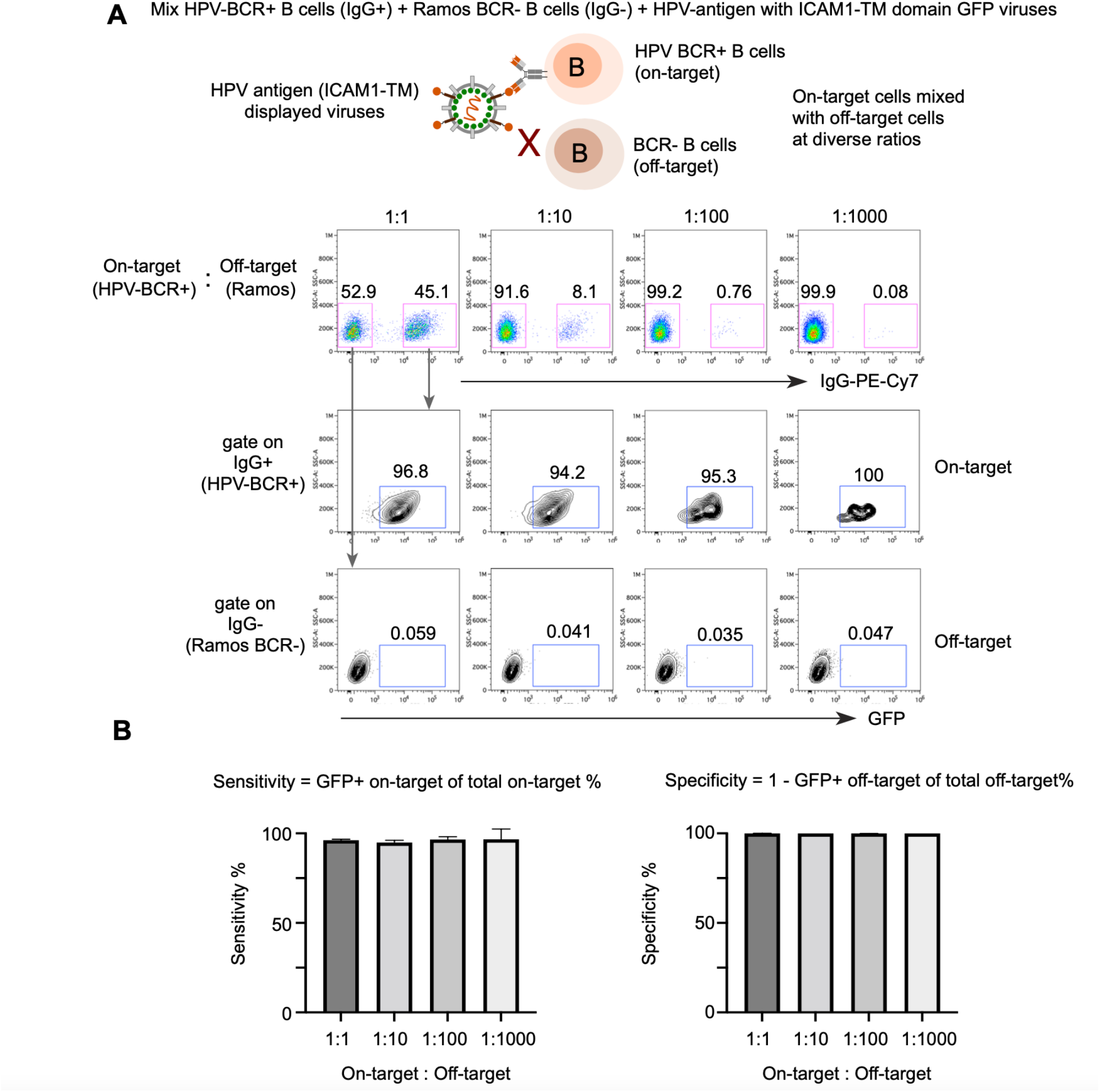
Specificity and sensitivity of ENTER in decoding interaction between BCR and antigens. A. Schematic view of experimental design (top) and flow cytometry analysis (bottom) of B cell mixing experiments. HPV-BCR+ B cells were mixed with BCR-B cells at different ratio. Mixed B cell population was incubated with GFP viruses displaying HPV epitope fused with ICAM1 TM domain and then subjected to flow cytometry. Representative flow cytometry plot showing mixing of two B cell population and GFP signal of B cells gated on IgG+ and IgG- population. B. Bar plots showing sensitivity (left) and specificity (right) of ENTER from Figure S3A.

**Figure S4.**
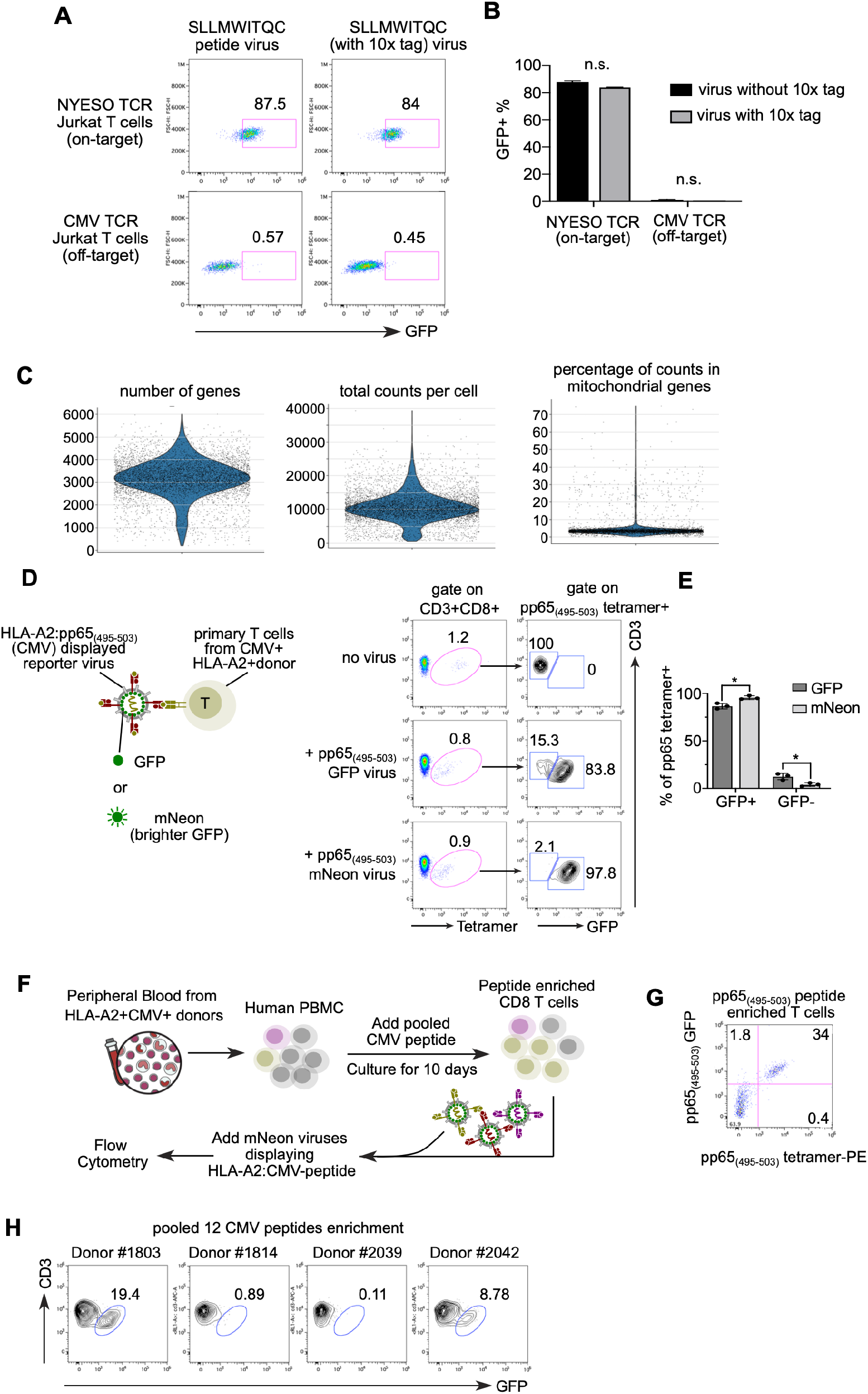
Optimize ENTER to detect rare antigen-specific primary human T cells. A. Flow cytometry analysis of GFP signal in T cells expressing TCRs (NYESO-TCR or CMV-TCR) upon incubation with nyeso_157-165_ antigen peptide displayed GFP viruses whose viral RNA are either intact or inserted with sequencing capture tag. B. Bar plot showing the percentage of GFP+ cells from Figure S4A. P value was calculated by unpaired t-test. n.s. P>0.05 C. Violin plots showing the quality control metrics of single cell gene expression by ENTER-seq. D. Schematic view of experimental design (left) and flow cytometry analysis (right) of tetramer and GFP signal in primary human T cells from donors with CMV infection. Donor isolated T cells were incubated with pp65_495-503_ displayed viruses carrying either GFP or mNeon fluorescence proteins and then stained with pp65_495-503_ tetramer and other antibodies followed by flow cytometry. We first gated on CD3+ CD8+ T cells to measure pp65-specific T cells using tetramer as a positive control, and then monitored GFP signals in pp65 tetramer+ T cells. E. Bar plot showing the percentage of GFP+ and GFP-cells among pp65 tetramer+ T cells from Figure S4D. F. Schematic view of experimental design. PBMCs were isolated from CMV seropositive HLA-A2+ donors and cultured with 10ug/mL CMV pp65 peptide for 10 days. Peptide stimulated PBMCs were incubated with pp65 antigen displayed mNeon viruses for 2 hours and then stained with antibodies followed by flow cytometry. G. Representative flow cytometry plot showing co-staining of pp65_495-503_ tetramer and pp65_495-503_ displayed mNeon viruses in pp65_495-503_ peptide enriched T cells. These T cells are gated on live CD8+ CD3+ T cells. H. Representative flow cytometry plot showing the percentage of GFP+ T cells (stained by 12 pooled HLA-A2:CMV-antigen mNeon viruses) from 4 different CMV seropositive donors upon pooled CMV peptide enrichment as in Figure S4F.

**Figure S5.**
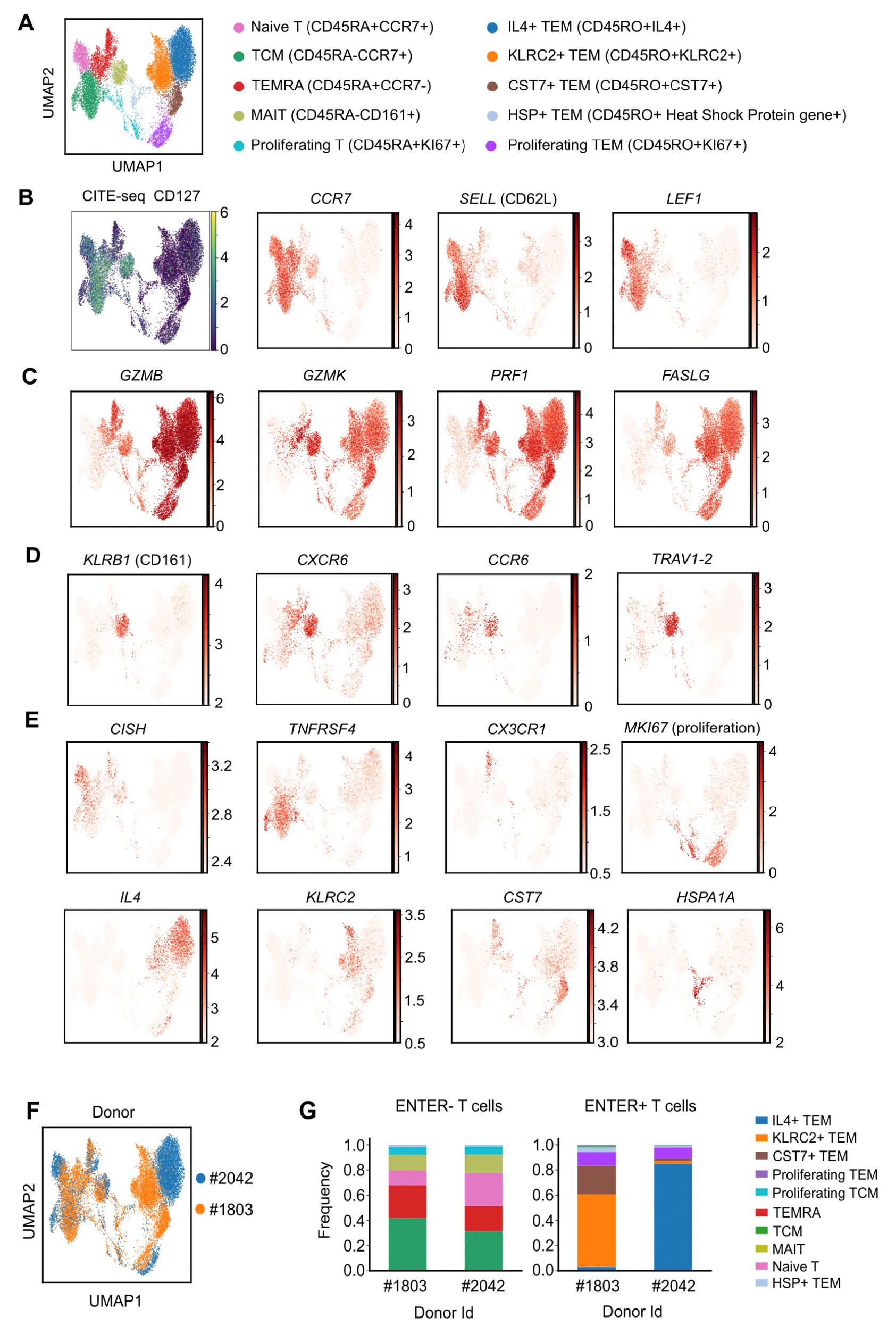
Characterization of T cell subsets by ENTER-seq. A. UMAP plot showing 10 clusters of human CD8 T cell subsets. B. UMAP plots showing amount of surface protein CD127 from CITE-seq, and expression of genes for naïve T cells (CCR7, SELL, and LEF1). C. UMAP plots showing gene expression of cytolytic molecules. D. UMAP plots showing gene expression of markers for MAIT cells. E. UMAP plots showing expression of marker genes for each subset/cluster. F. UMAP plot showing donor origin. G. Fraction of T cell subsets from 10 clusters in CMV antigen-specific T cells (ENTER+) and bystander T cells (ENTER-), separated by donor origin.

**Figure S6.**
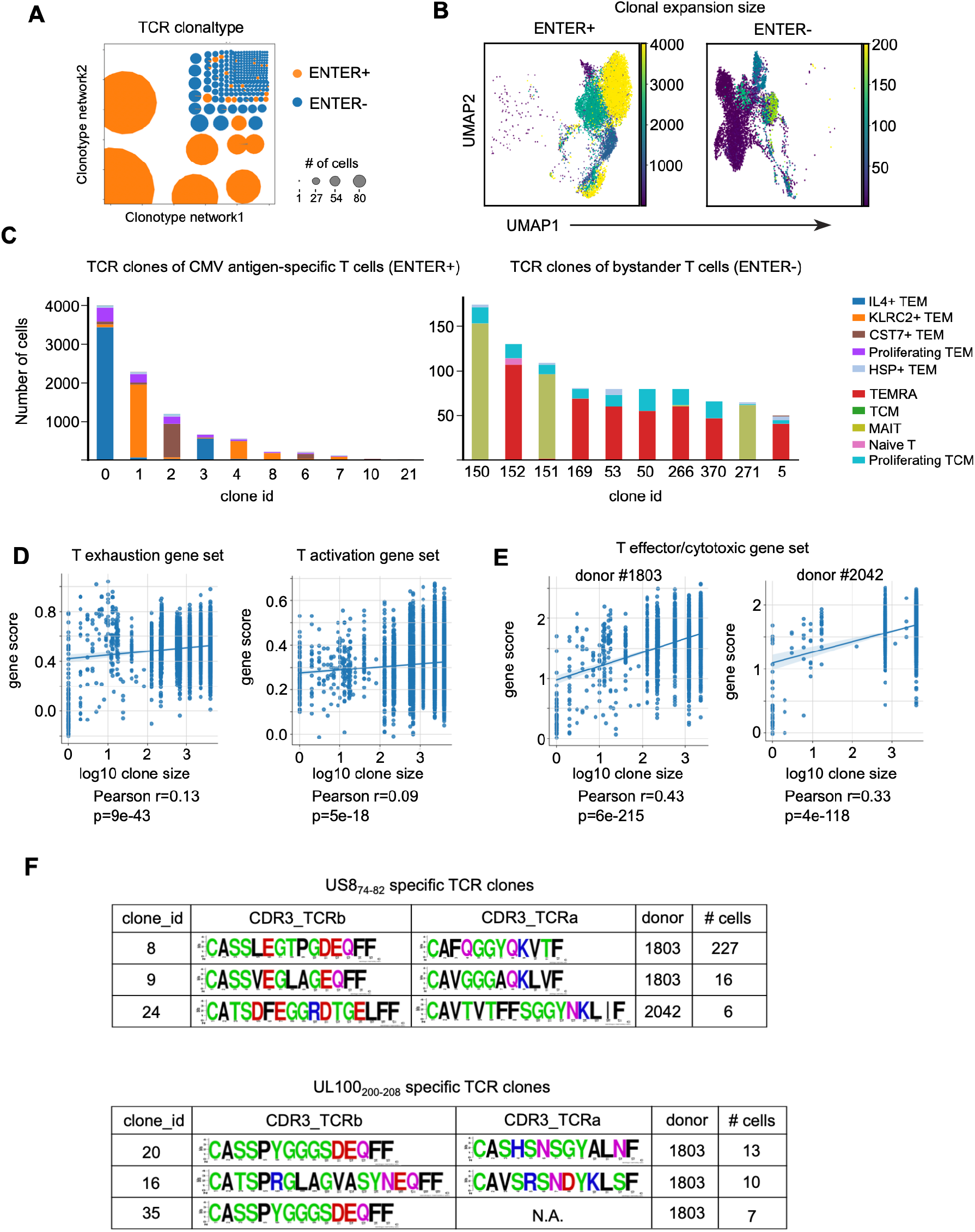
TCR clonotype analysis from ENTER-seq. A. TCR clonotypes colored by ENTER+ (CMV antigen-specific T cells) and ENTER- (bystander T cells). The size of circle represents the number of cells per clonotype. B. UMAP plots showing the clonal expansion size of CMV antigen-specific T cells (ENTER+) and bystander T cells (ENTER-). C. The number of ENTER+ or ENTER-T cells in top 10 TCR clones, colored by the T cell subsets from 10 clusters and separated by TCR clone id. D. Scatter plots showing the correlation of clonal expansion size with exhaustion gene score or activation gene score. E. Scatter plots showing the correlation of clonal expansion size with effector gene score in each donor. F. Summary tables of US8_74-82_ - and UL100_200-208_ - specific TCR clones. Coefficient r and p values in Figure S6D-E are calculated by Pearson correlation.

**Table S1.**
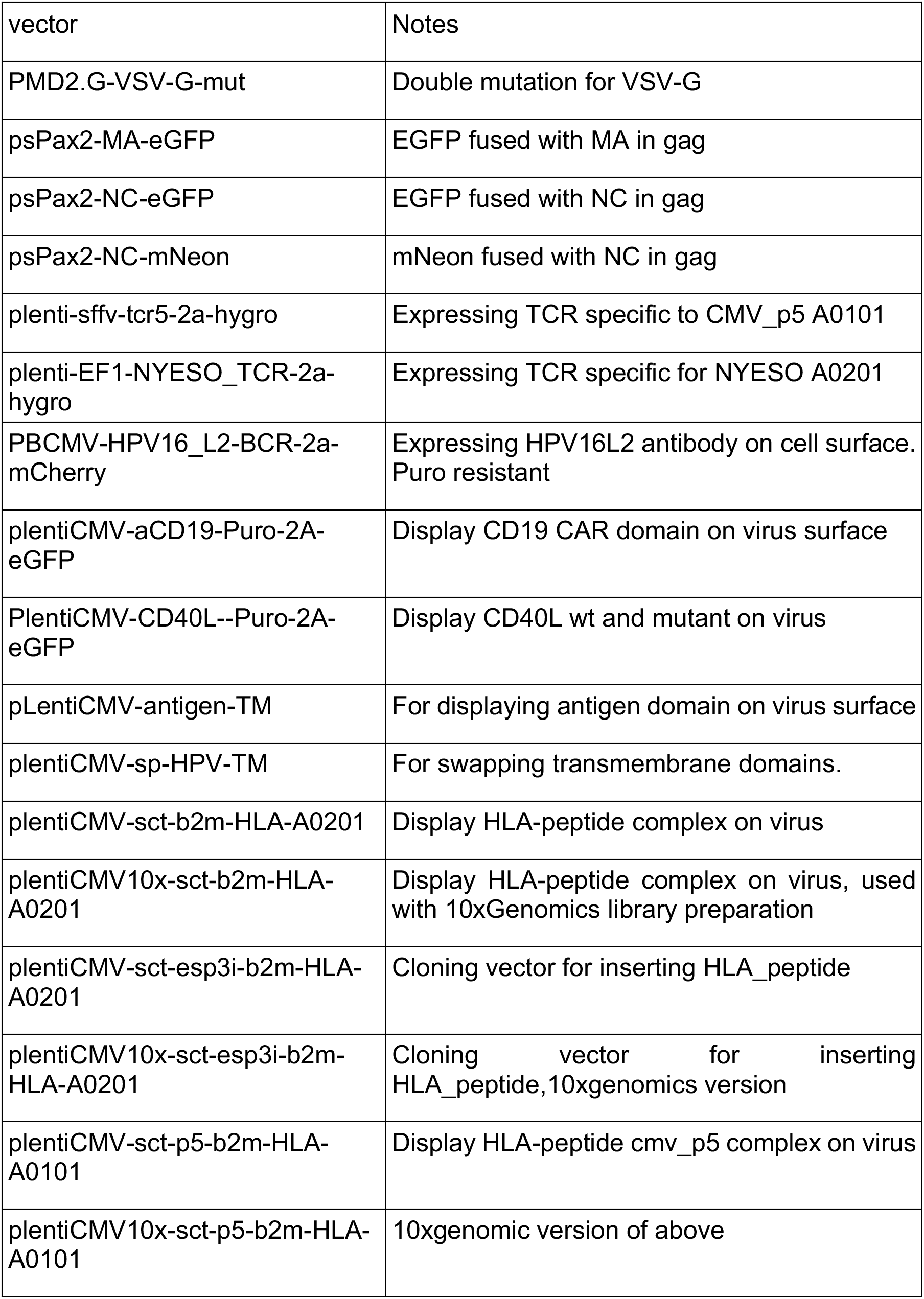
List of vectors made.

**Table S2.**
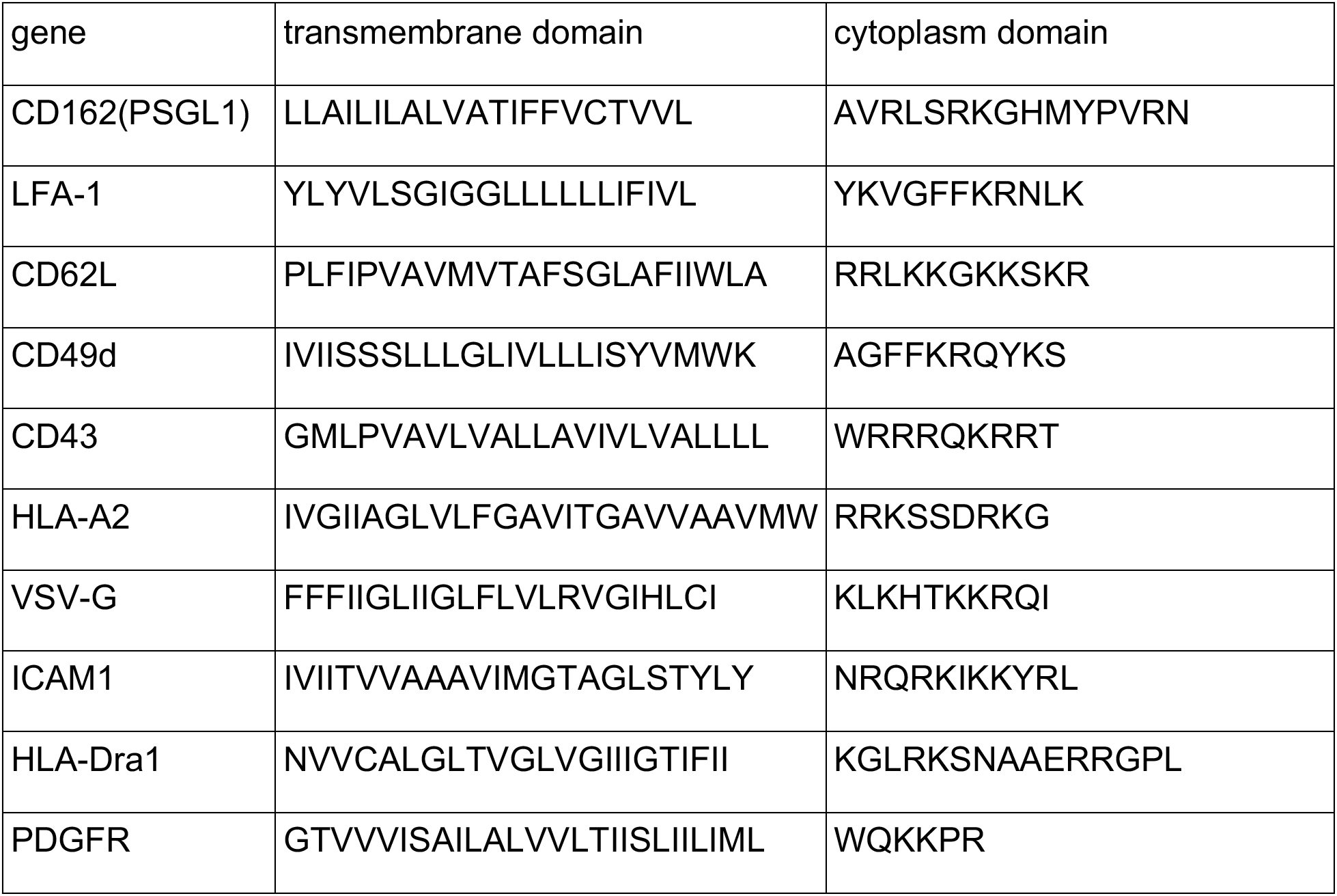
List of TM domain sequence.

**Table S3.**
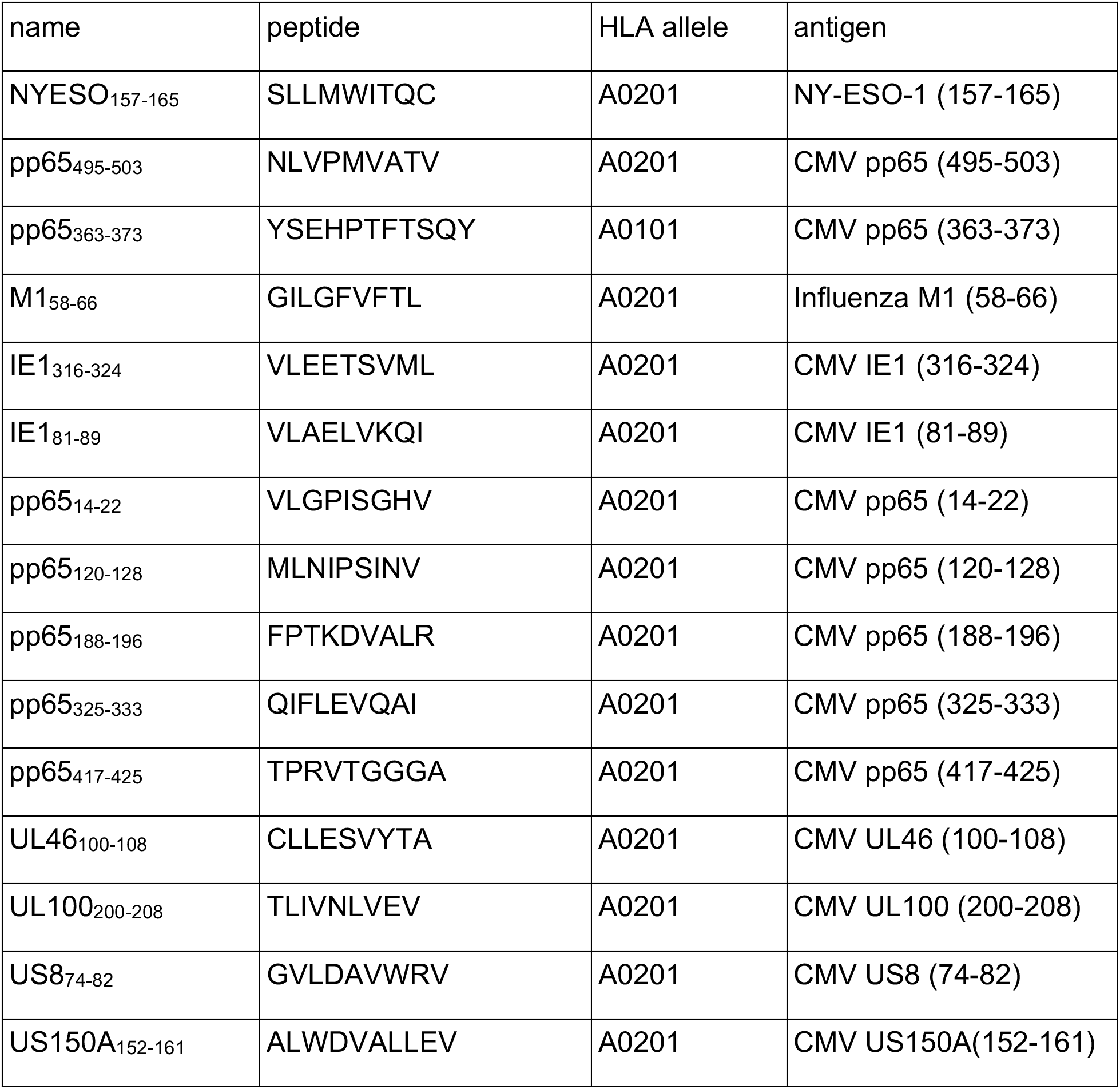
List of HLA peptide sequence.

**Table S4.**
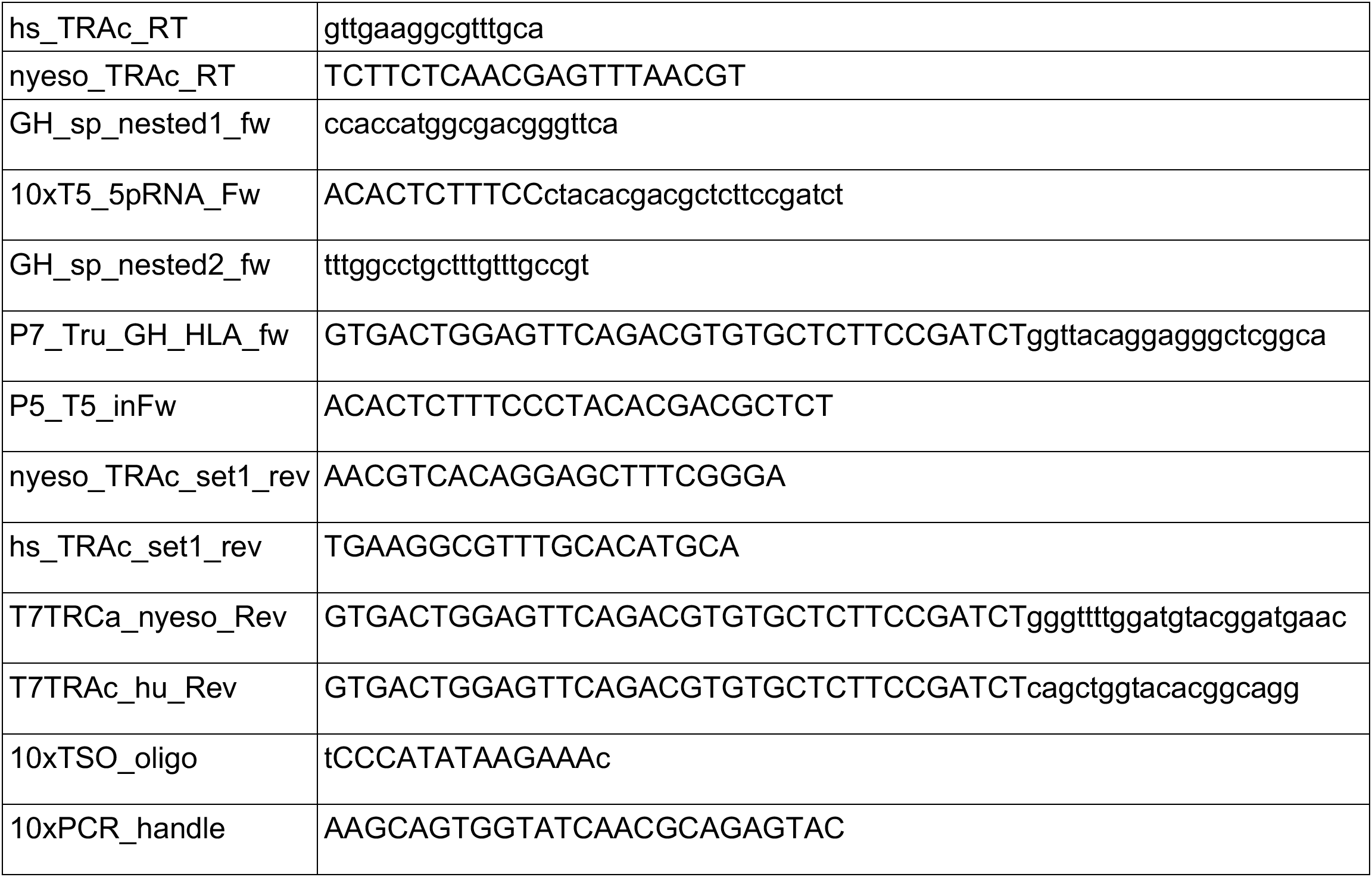
DNA Oligo sequences.

